# Mitochondria-derived Vesicles mediate Mito-nuclear Trafficking of Cargo

**DOI:** 10.1101/2024.05.04.592299

**Authors:** Thejaswitha Rajeev, Dipak Shil, Swagato Bhattacharjee, Lariza Ramesh, Victor Samuel, Nidhi Nair, Areeba Marib, Mrithulabhashini Jayakumar, Sai Gayathri, Arun Surendran, John B Johnson, Ananthalakshmy Sundararaman

## Abstract

Mito-nuclear communication is key to cellular homeostasis. Metabolites from TCA cycle, calcium, ROS and other small molecules are known to mediate retrograde signalling of mitochondria to the nucleus. Recently, the direct transfer of mitochondrial proteins to the nucleus has been described. However, the mechanism for direct transfer of mitochondrial protein complexes to the nucleus is not understood. In this study, we demonstrate the transit of mitochondrial proteins to the nucleus through mitochondria-derived vesicles (MDVs) which represents a novel intracellular trafficking route. Following an unbiased proteomic approach to detecting cargo proteins in cardiac MDVs, we screen for the presence of MDV cargo in the nucleus. Pyruvate dehydrogenase complex (PDH) and Cytochrome c oxidase subunit IV isoform 1 (COX4I1) are packaged into MDVs in cardiomyocytes but only PDH is targeted to the nucleus in this cell type demonstrating that a subset of cargo-selective MDVs fuse with the nucleus. Further, several mitochondrial stresses that are known to increase MDV generation triage these MDVs to other cellular destinations thereby reducing the nuclear pool of PDH. The transit of PDH through MDVs to the nucleus is a basal physiological process in cardiomyocytes. Mito-nuclear trafficking thus represents a physiological pathway in cardiac cells to enable cargo shuttle from mitochondria to the nucleus.

## Introduction

Cardiomyocytes rely heavily on mitochondria for energy through oxidative phosphorylation which contributes to 90% of cellular ATP levels [1]. Mitochondrial dysfunction contributes to various cardiovascular diseases. Hence there exists various mitochondrial quality control pathways to ensure a healthy mitochondrial network [2]. One such quality control mechanism discovered recently is through mitochondria-derived vesicles (MDVs) that transport oxidatively damaged proteins to lysosomes for degradation [3]. While MDV biology is not yet explored in depth in cardiac cells, they are present in a basal physiological state in this system [4]. Interestingly, fission-fusion cycle is extremely rare in cardiomyocytes [5] which is an important quality control mechanism in other cell types. Therefore, we hypothesise that cardiac cells would be highly reliant on MDVs to maintain mitochondrial homeostasis and for inter-organellar communication. Our study therefore aims to understand MDV biogenesis, its trafficking, particularly to the nucleus and its relevance to the cardiac system.

Mitochondria-derived vesicles are cargo selective vesicles of 70-150 nm in size that originate from mitochondria. They can be single or double membrane structures that can target cargo to cell destinations like the peroxisome and lysosome [3, 6]. MDV generation and targeting is aided by various proteins including Vps35, PINK1/PARKIN, syntaxin 17-SNARE complex, Sorting nexin 9-Rab 9, Rab 7, and Tollip [7–13]. Previous studies have shown that MDVs are formed independent of the fission protein, Drp1 [6, 10]. But, in the case of the widely studied TOM20^+^ MDVs, MIRO1/2 and Drp1 are required for MDV formation [14]. MDVs thus generated play a variety of roles including peroxisome biogenesis, mitochondrial quality control, mitochondrial antigen presentation, bacterial killing within phagosomes and in the regeneration of ATP deficient mitochondria [3, 6, 10, 15, 16]. However, MDVs have not been implicated in mito-nuclear cargo transit which we explore here.

Mitochondria in eukaryotes evolved from an α-proteobacterial ancestor capable of secreting outer membrane vesicles (OMV) for inter-colony communication and interaction with host cells [17–19]. As MDVs are speculated to be evolutionarily descended or closely related to bacterial OMVs we fathom that inter-organellar communication, particularly between the endosymbiont and host nuclei might be a key physiological role for MDVs. We therefore hypothesised that MDVs mediate mito-nuclear protein transit. Several mitochondrial proteins are known to have moonlighting functions in the nucleus. For example, Nuclear factor erythroid 2-related factor 2 (NRF2), CLOCK-1 (CLK-1), fumarase, Lonp1 and TCA cycle enzymes are mitochondrial proteins also found to function in the nucleus in different systems [20–24]. In mouse embryo, pyruvate carboxylase (PCB), citrate synthase (CS), aconitase 2 (ACO2), isocitrate dehydrogenase 3 (IDH3A) and pyruvate dehydrogenase (PDHE1) are required in the nucleus for early zygotic genome activation (ZGA) [25]. The same enzymes have been found to play a role in somatic cell reprogramming as well [26]. In the only study exploring moonlighting functions of TCA enzymes in the cardiac system, malate dehydrogenase 2 (MDH2), pyruvate dehydrogenase subunit 1 (PDH-E1) and isocitrate dehydrogenase 2 (IDH-2) were found in the nucleus of hiPSC-derived cardiomyocytes upon doxorubicin-mediated cardiotoxicity [27]. A well studied nuclear translocated mitochondrial protein is pyruvate dehydrogenase (PDH) [28, 29] which is also independently characterised as an MDV cargo [3, 30].

Pyruvate dehydrogenase complex (PDC) is a ∼10 MDa, multi-subunit complex of diameter 45 nm, linking glycolysis and TCA cycle through acetyl CoA generation. The subunits are PDHE1 (pyruvate dehydrogenase), PDHE2/DLAT (dihydrolipoamide acetyltransferase), PDHE3 (dihydrolipoamide dehydrogenase), and E3BP (E3 binding protein) [29, 31, 32]. It has been shown that the nucleus accumulates only the mitochondria localization signal (MLS) cleaved mature form of PDC subunits and since the peptidase to process this is available only in the mitochondria it stands to reason that PDH reaches the nucleus from the mitochondria. Nuclear translocation of PDC occurs when cells are exposed to proliferative stimuli or mitochondrial stresses like hypoxia [28, 29]. But prolonged hypoxia can lead to decreased nuclear PDH [33]. Within the nucleus, PDH is believed to regulate gene expression through histone acetylation [26, 28, 29] as seen in the regulation of the expression of Sterol regulatory element-binding transcription factor (SREBF) by nuclear PDC in prostate cancer [34]. So, nuclear PDC has several cell type-specific roles like cardioprotective response to cardiotoxicity, tumour progression, epigenetic reprogramming after T cell activation [27, 34, 35].

Here, we hope to address the question of the entry of pyruvate dehydrogenase complex which cannot freely diffuse nor pass through the nuclear pore complex (NPC) [29]. We propose that packaging of either the pyruvate dehydrogenase complex or their individual subunits into MDVs as a means of mito-nuclear trafficking. MDVs labelled with PDHE2/E3bp are reported [36] but these are believed to exclude PDHE1 [8]. On the other hand, some papers show PDHE1^+^ MDVs but their cellular destinations are not explored [10, 37]. MDV transit and corresponding changes to nuclear pool of cargo proteins have not been studied in tandem. It is reported that PDH^+^ MDV number increases in response to xanthine oxidase as well as antimycin-A treatment [4, 30]. PDH^+^ MDV biogenesis also increases in cardiotoxicity after chemotherapy [4] but in these cases the cellular destination of PDH^+^ MDVs remain unexplored. Interestingly, PDH^+^ MDVs can triage to different cell destinations based on cellular context, for example Parkin dependent MDVs to lysosomes and Parkin independent MDVs to endosomes and extracellular vesicles [8, 9, 11]. In this study we unravel the cellular context under which PDH in MDVs could be directed to the nucleus thereby describing an unknown trafficking route for MDVs.

## Results

### Cardiac MDV proteome is enriched for mitochondrial proteins involved in TCA cycle and electron transport chain

We initially took an unbiased proteomic approach to generate MDV cargo list from rat hearts through *in vitro* budding assay. This approach has been used previously [30, 36, 38] but our budding reactions additionally incorporate N-ethyl maleimide (NEM) [39]and we further analyse cargos based on their NEM sensitivity. Figure 1A shows the steps in our cell free MDV budding assay. A detailed protocol is available from our group in Nair et al., 2024 [40].

**Figure 1:**
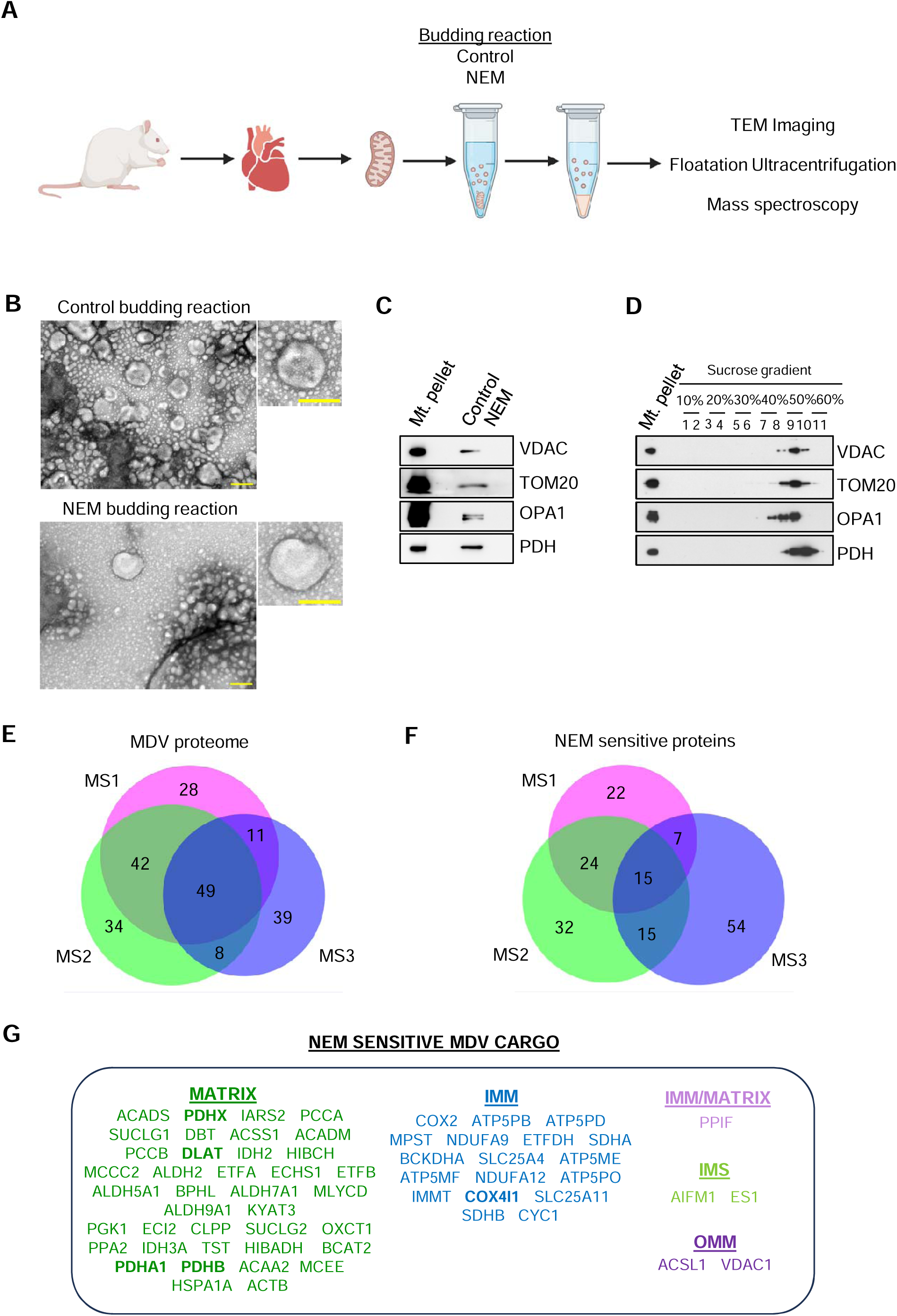
Cardiac MDV proteome was enriched for mitochondrial proteins involved in TCA cycle and electron transport chain. (A) Schematic of the MDV isolation protocol, created with BioRender.com. Mitochondria isolated from rat heart were budded in a budding reaction buffer (composition is as described in methods) for 2h at 37°C. In parallel reactions, the budding was performed in the presence or absence of vesicular transport inhibitor, N-ethylmaleimide (NEM: 20mM). The mitochondria was pelleted and the supernatant containing the MDVs was used for downstream experiments. (B) TEM imaging of the budding reactions performed in the presence or absence of NEM showed vesicular structures. Scale bar shows 100 nm. (C) Immunoblot of the budding reactions generated in the presence or absence of NEM. Equal volumes of the budding reactions was loaded and were probed for outer mitochondrial membrane proteins, Voltage-dependent anion channel (VDAC) and Translocase of outer mitochondrial membrane 20 (TOM20); inner mitochondrial membrane and intermembrane space protein, Optic atrophy 1 (OPA1) and mitochondrial matrix protein, Pyruvate dehydrogenase E1 (PDHE1). Mt. pellet: mitochondria pellet. (D) Immunoblot of fractions collected after sucrose density gradient (10-60%) ultracentrifugation of budding reaction. Blot was probed for VDAC, TOM20, OPA1 and PDHE1. Mt. pellet: mitochondria pellet. (E) Venn diagram showing the overlap of the mitochondrial proteins detected in the three MS runs. (F) Venn diagram showing the overlap of the NEM sensitive proteins. (G) List of the NEM sensitive proteins. These proteins appeared at least twice in three independent MS runs.

To confirm that the control and NEM budding reactions contained vesicles, the reactions were visualised using TEM. We observed vesicular structures ranging from 70 to 150 nm in size which is within the size range of MDVs [6]. There were less vesicular structures in the NEM budding reaction as NEM is expected to reduce the budding of the vesicles (Figure 1B).

To further confirm our protocol, we immunoblotted the budding reaction with some of the previously reported MDV cargo like VDAC, Tom20, Opa1 and PDH [3, 11, 30, 38]. We confirmed that all these cargo proteins are incorporated into MDVs in an NEM sensitive manner (Figure 1C). For further validation of the packaging of these proteins in vesicles, we floated the control budding reaction in a sucrose gradient by ultracentrifugation. We observed that the matrix protein, PDH, was also present in the denser sucrose fractions compared to all other proteins from other compartments (Figure 1D). Together our TEM, immunoblotting of the budding reactions and floatation gradients validate our in vitro approach to generation of MDVs from rat heart.

Having validated the cell free MDV budding, we proceeded to identify the cargo packaged within these MDVs by mass spectrometry (Supplementary Data S1). We used multiple sources including MitoCarta 3.0 and Integrated Mitochondrial Protein Index (IMPI) database to confirm the mitochondrial localization and sublocalization of the detected proteins. Of the total 453 proteins detected in the three independent MS analyses of rat cardiac MDV proteome, 211 proteins were mitochondrial proteins. Out of the 211 mitochondrial proteins only those proteins that appeared twice in the three MS runs were analysed further as MDV cargo. There were 110 mitochondrial proteins primarily of matrix and inner mitochondrial origin in our MDV proteome (Figure S1A). This preponderance of IMM and matrix proteins in the in vitro generated MDVs is also reported previously [36]. Pathway analysis indicated that these proteins belong mostly to the group of TCA cycle enzymes and electron transport chain proteins (Figures S1C and S1D).

We next considered NEM-sensitive proteins (present in the control reaction but absent in the NEM containing reaction) as a more stringent list of MDV cargo. This eliminates any artifactually present proteins in the supernatant post-budding. When NEM sensitivity was used as cut-off 61 proteins were identified as MDV cargo in at least two out of the three MS runs (Figure 1G). These were also predominantly matrix and inner mitochondrial membrane proteins playing roles in TCA cycle and electron transport chain (Figure S1B).

In our NEM sensitive list of MDV cargo, we noted multiple subunits of the PDC complex like PDHA1 (PDHE1α), PDHB (PDHE1β), DLAT (PDHE2) and PDHX (PDHE3 binding protein) (Figure 1G) which have previously been used to mark matrix positive MDVs in the field [8, 10]. Additionally, we noticed both subunits of SDH and components of the complex IV of the respiratory electron transport chain. We validated the new hits like COX4I and SDH as being incorporated into MDVs in an NEM sensitive manner *in vitro* (Figure S1E) and further demonstrated the presence of COX4I1 and SDH in MDVs floated on sucrose gradients (Figure S1F).

Next, we validated the formation of MDVs in our cells of interest, H9C2 cardiomyoblasts. PDH^+^ MDVs were already known to exist basally in H9C2 cardiomyoblasts and were upregulated in the presence of an oxidative stress induced by antimycin A [4]. We observed COX4^+^PDH^-^, PDH^+^COX4^-^ and COX4^+^PDH^+^ dual positive MDVs basally and found that these MDVs were antimycin sensitive (Figures S1 G-K).

Together our NEM sensitive MDV cargo part list, while likely incomplete, adds to the set of high confidence MDV cargo that can be used to understand MDV biology in the cardiac system.

### PDH, but not COX4, is found in the nucleus of rat cardiac cells

To examine if any of the mitochondrial proteins that we identified as MDV cargo were localized to the nucleus, we performed an initial screening in the nuclei of HEK293T cells.

We detected mitochondrial proteins from various sublocalizations in the nuclear isolates of HEK293T (Figures 2A and S2A). We observed the outer mitochondrial membrane proteins (VDAC and TOM20) in the nucleus. We also observed COX4, the newly identified MDV cargo in the nucleus. COX4 is a subunit of the Complex IV of the ETC. We did not detect the other subunits of Complex IV, COX1 and COX10 in the nucleus suggesting that not the entire Complex IV but subassemblies may be transiting to the nucleus. Prohibitin and NDUFS1 are inner membrane proteins detected in the nucleus in our system. While Prohibitin is already known to be present in the nucleus [41], the other proteins are not known to be present in the nucleus. Several of the other inner mitochondrial proteins that were part of or closely associated with the ETC, including RISP, ACP, SDH and cytochrome c were absent in the nucleus. Also, inner membrane protein involved in membrane fission Opa1, was absent in the nucleus.

**Figure 2:**
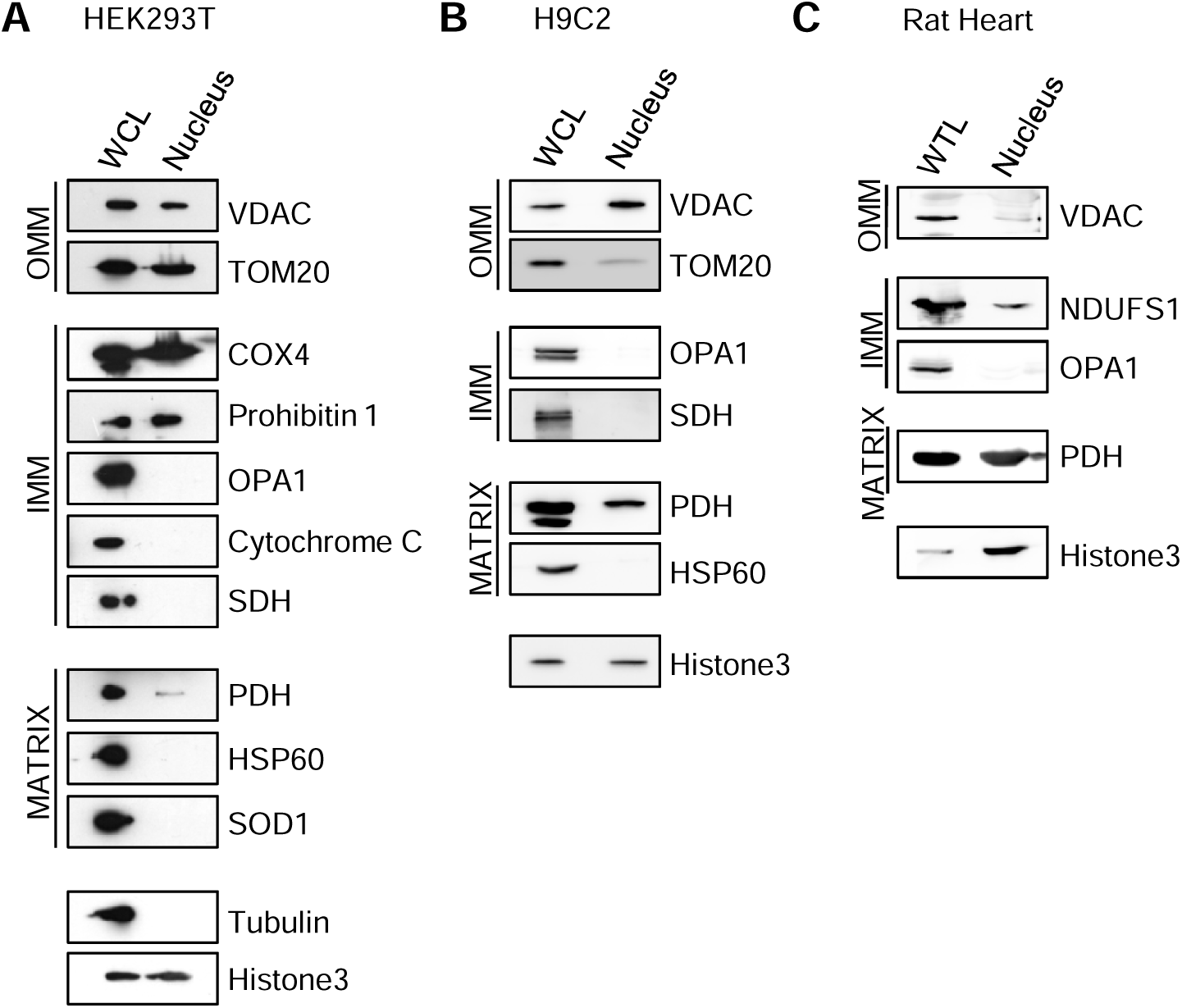
PDH, but not COX4, is found in the nucleus of rat cardiac cells. (A) Immunoblot of the nuclei isolated from HEK293T. The blot was probed for mitochondrial proteins(VDAC, TOM20, COX4, Prohibitin 1, OPA1, Cytochrome, SDH, PDHE1, HSP60, SOD1); cytoplasmic marker, Tubulin and nuclear marker, Histone 3. WCL: whole cell lysate. (B) Immunoblot of the nuclei isolated from H9C2 cardiomyoblasts. The blot was probed for mitochondrial proteins(VDAC, TOM20, OPA1, SDH, PDHE1, HSP60) and Histone 3. WCL: whole cell lysate. (C) Immunoblot of the nuclei isolated from rat heart. The blot was probed for mitochondrial proteins(VDAC, NDUFS1, OPA1, PDHE1) and Histone 3. WTL: whole tissue lysate

We observed trace amounts of PDH in the nuclei of HEK293T. PDH was previously reported in the nucleus of other cell types [25, 28, 42, 43]. Other matrix proteins like SOD1 and HSP60 were not detected in the nucleus. Of the screened proteins detected in the nucleus, PDH, TOM20, VDAC and COX4 were MDV cargo validated by us and other groups.

Next, we investigated the presence of mitochondrial proteins in the nucleus of H9C2 cardiomyoblasts and rat heart nuclei. We observed VDAC in the nuclei from both H9C2 and rat heart but detected TOM20 only in H9C2 cardiomyoblasts. We did not detect COX4 or any of the other Complex IV subunits (COX1 and COX10) in the nuclei of either H9C2 cardiomyoblasts or rat heart. But PDH was detected in the nucleus in cardiac cells (Figures 2B-C, S2B-C). In H9C2 cardiomyoblasts overexpressing PDH mCherry, we observed PDH^+^ MitoTracker ^negative^ vesicular structures near the mitochondria before their fusion with the nucleus to release PDH in the nucleoplasm(Figure S2E, Supplementary movie Data S2). We also observed that in endothelial cells, PDH but not COX4 is present in the nucleus (Figure S2D). Together, we confirm that nuclear transit of proteins including PDH and COX4 are cell type specific and PDH but not COX4 is present in the nucleus of cardiac and endothelial cells.

It is noteworthy that not all of the screened MDV cargo gets delivered to the nucleus as observed in the cases of COX4, OPA1 and SDH. This highlights the specificity of mito-nuclear protein transit. While the screen does not provide evidence for the transit being MDV-mediated, lack of transit of some proteins and the cell type specificity in others ensure that this is not an artifactual co-purification of some mitochondria with the nuclear pool. We further explore PDH transit in detail as it is present in the nuclei of cardiomyoblast cells and the rat heart tissue.

### Actin and tubulin depolymerization leads to decreased PDH^+^ MDVs

It is known that TOM20^+^ MDVs are trafficked to endolysosomes in a microtubule dependent manner with the aid of MIRO1/2 and DRP1 [14]. However, other cytoskeletal elements like actin which is known to control vesicle formation has not been studied in the context of MDV biogenesis and trafficking [44, 45]. We wanted to explore the role of actin and tubulin in PDH^+^MDV biogenesis in cardiomyocytes.

We had previously observed that antimycin A upregulates the number of PDH^+^ MDVs formed. We sought to investigate the potential impact of actin or tubulin depolymerization on antimycin-driven mitochondrial-derived vesicles (MDVs). First, we had confirmed that the actin and tubulin depolymerized at the concentrations (1µM and 10 µM respectively) used in our study (Figures S3A and S3B). We observed a significant drop in the number of PDH^+^

MDVs formed in the presence of latrunculin B (Figures 3A and 3B) or nocodazole (Figures 3C and 3D). This is evidence that cytoskeletal elements like actin and tubulin play a role in PDH^+^ MDV biogenesis. To uncouple MDV trafficking and nuclear fusion events from budding we undertook an *in vitro* reconstitution approach.

**Figure 3:**
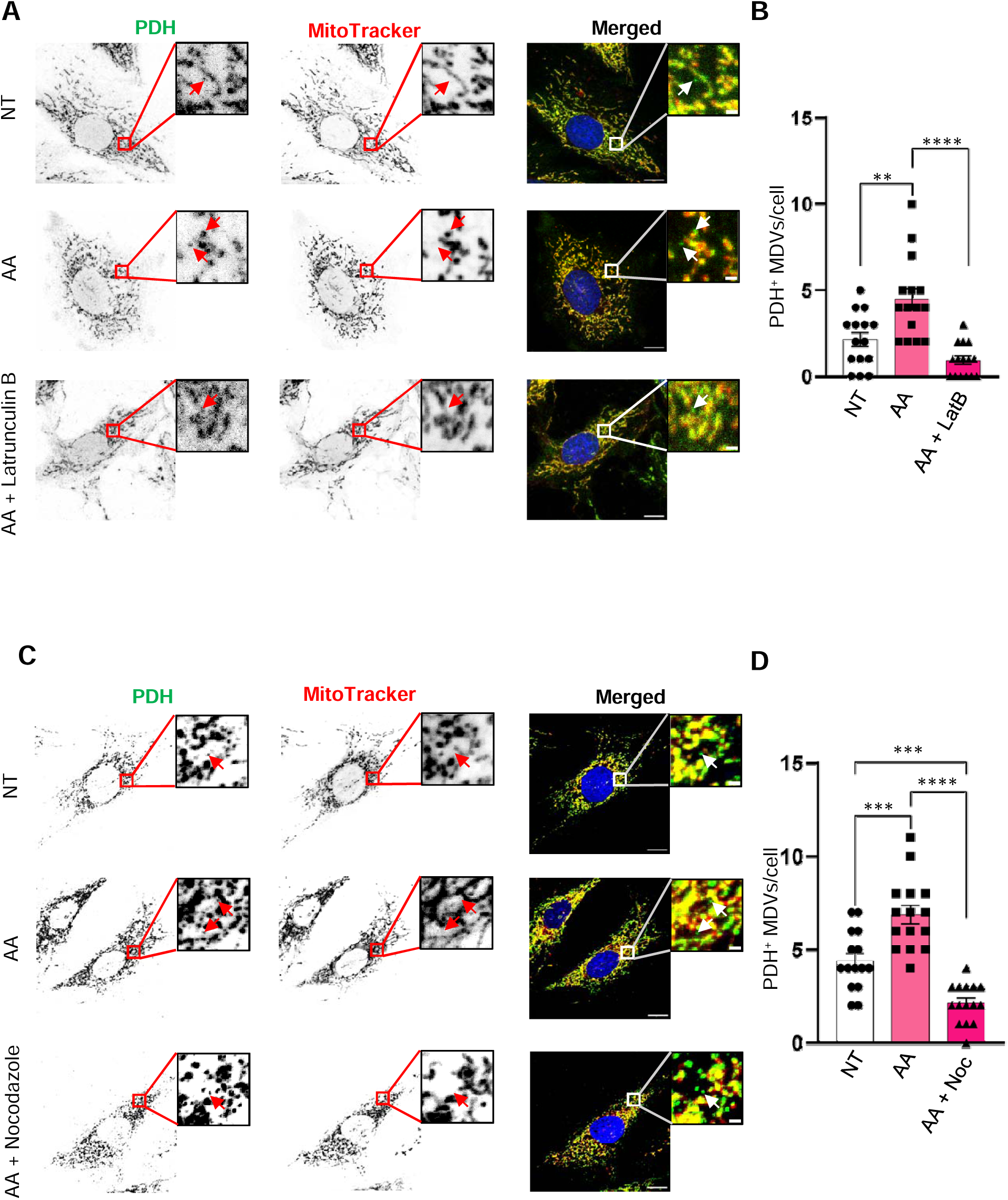
Actin and tubulin depolymerization leads to decreased PDH^+^ MDVs. (A) H9C2 rat cardiomyoblasts were pre-treated with MitoTracker (500nM) for 30 min. Subsequently the cells were treated with vehicle (NT), AA (20µM) and Lat B (1µM) with AA. The cells were fixed after 15 min with 100% methanol at −20°C. The cells were immunolabeled for PDH E1-α (green) and the nucleus was stained with DAPI (blue). MitoTracker appears red. The arrowhead indicates MDVs. Scale bar:10 µm (cell), 1 µm (inset). (B) Quantification of PDH^+^ MDVs (N=3, 5 cells/condition). One-way ANOVA followed by Bonferroni post-hoc analysis, ****p-value< 0.0001, **p-value< 0.01. Error bars, SEM. (B) H9C2 rat cardiomyoblasts were pre-treated with MitoTracker (500nM) for 30 min. Subsequently the cells were treated with vehicle (NT), AA (20µM) and Noc (10µM) with AA. The cells were fixed after 30 min with 100% methanol at −20°C. The cells were immunolabeled for PDH E1-α (green) and the nucleus was stained with DAPI (blue). MitoTracker appears red. The arrowhead indicates MDVs. Scale bar:10 µm (cell), 1 µm (inset). (D) Quantification of PDH^+^ MDVs (N=3, 5 cells/condition). One-way ANOVA followed by Bonferroni post-hoc analysis, ****p-value< 0.0001, ***p-value< 0.001. Error bars, SEM.

### *In vitro* reconstitution of mito-nuclear trafficking shows that PDH delivery to the nucleus is NEM sensitive

To determine the mechanism of transit of mitochondrial proteins to the nuclei, we incubated rat cardiac MDVs with rat heart nuclei as shown in the graphical representation (Figure 4A). GTP (0.5 µM) was added to all the reconstitution reactions. The nuclear purity of our isolation could be noted by the absence of OPA1 and COX4 in the nuclear isolates as observed previously. We observed increased nuclear PDH in rat nuclei incubated with MDVs (Figures 4B and 4C) providing direct evidence for the delivery of PDH by MDVs to the nucleus. By addition of NEM to the cell free reconstitution reaction, we inhibited trafficking proteins required for the fusion of membranes. We noted that PDH entry into the nucleus was NEM sensitive. This proved that the fusion of MDVs with nucleus and PDH delivery required NEM sensitive proteins. We also observed that some of the other MDV cargo like NDUFS1 and VDAC did not appear to increase upon MDV addition or further reduce with NEM (Figure S4A). Therefore, these two proteins might enter the nucleus without MDV mediated shuttling.

**Figure 4:**
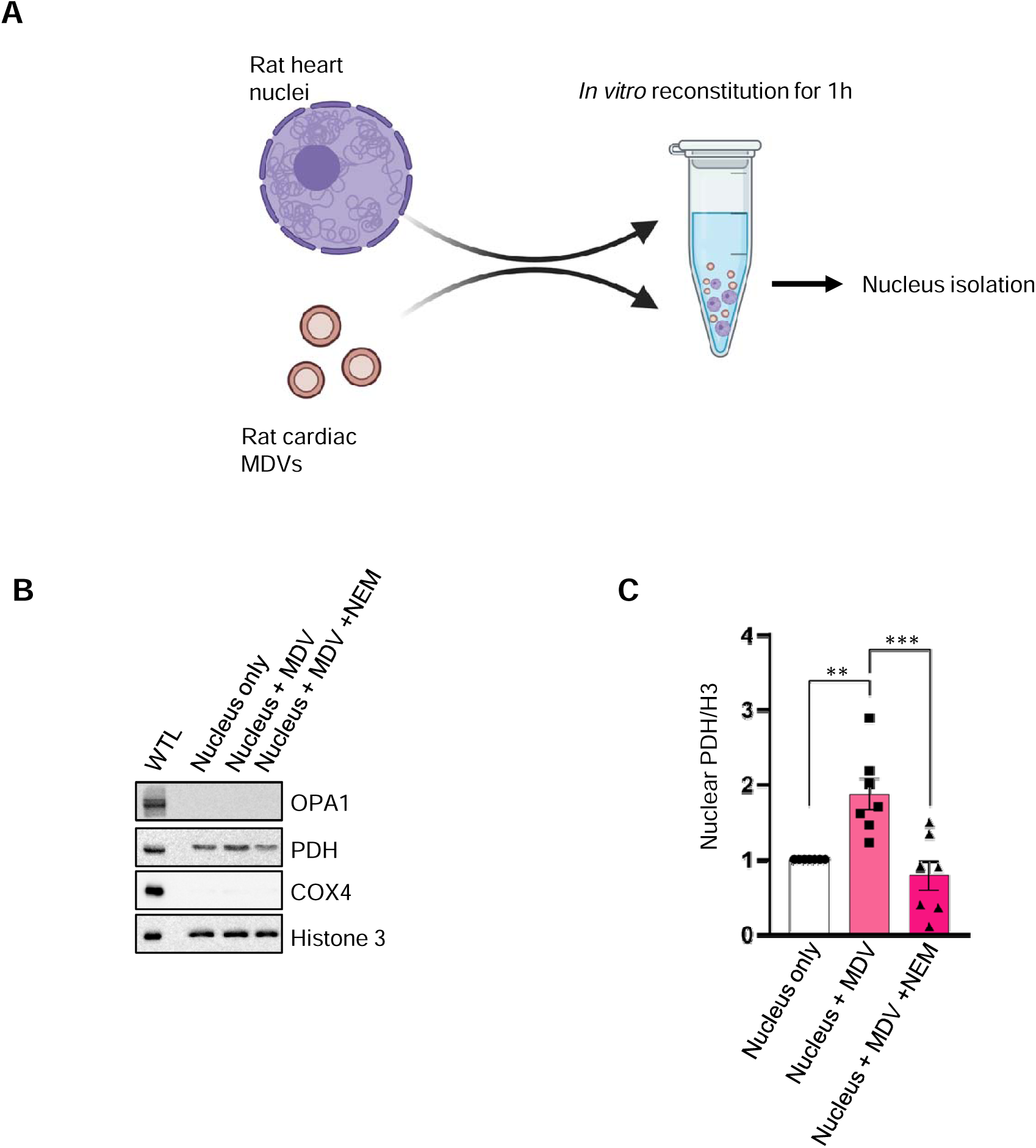
PDH delivery to the nucleus is NEM sensitive. (A) Schematic of the *in vitro* reconstitution between rat cardiac MDVs and rat nuclei, created with BioRender.com. Cardiac MDVs and nuclei were incubated together *in vitro* for 1 hour at 37°C. Three parallel reconstitution conditions were set up: nucleus only; nucleus and cardiac MDVs; and nucleus and MDVs in the presence of NEM (20mM). The nuclei were subsequently isolated by ultracentrifugation. (B) Immunoblot of the nuclei isolated after reconstitution as described in (A). The whole tissue lysate (WTL) was prepared from rat heart tissue using Laemmli buffer followed by sonication. The blot was probed OPA1, PDHE1, COX4 and Histone 3 (C) Quantification of nuclear PDH after reconstitution. Nuclear PDH was normalized to Histone 3. One-way ANOVA followed by Bonferroni post-hoc analysis, ***p-value< 0.001, **p-value< 0.01. Error bars, SEM.

### Mitochondrial stress represses PDH transit to the nucleus

As our previous results indicate a transit of PDH to the nucleus in cardiac cells, we wanted to understand the physiological and pathophysiological conditions under which this transit is modulated. When cells are grown in galactose containing medium, they mostly rely on mitochondrial function through oxidative phosphorylation to generate ATP [46, 47]. When we probed for PDH in the nuclear isolates of cells grown in the presence of glucose and compared it with 20 mM galactose containing media (with a low-5 mM glucose) we found that the nuclear PDH levels increase almost 1.5 folds in galactose containing medium (Figures 5A and 5B). Concomitantly we noted that the total cellular PDH levels remain unchanged in H9C2 cells when cultured for 24 hours in galactose containing media (Figures S5A and S5B). This suggests that PDH is transiting from the mitochondria to the nucleus under these conditions. Previously, it was shown that PDH and other TCA cycle enzymes transit to the nucleus in a pyruvate dependent manner during ZGA [25]. We assessed the role of pyruvate in nuclear PDH transit in H9C2 cardiomyoblasts. However, in our system no significant change was observed in nuclear PDH levels in the absence or presence of pyruvate in cells cultured in either glucose or galactose medium (Figures 5A and 5B). Again, we confirmed that pyruvate did not affect the total levels of PDH in the cells (Figures S5A and S5B).

**Figure 5:**
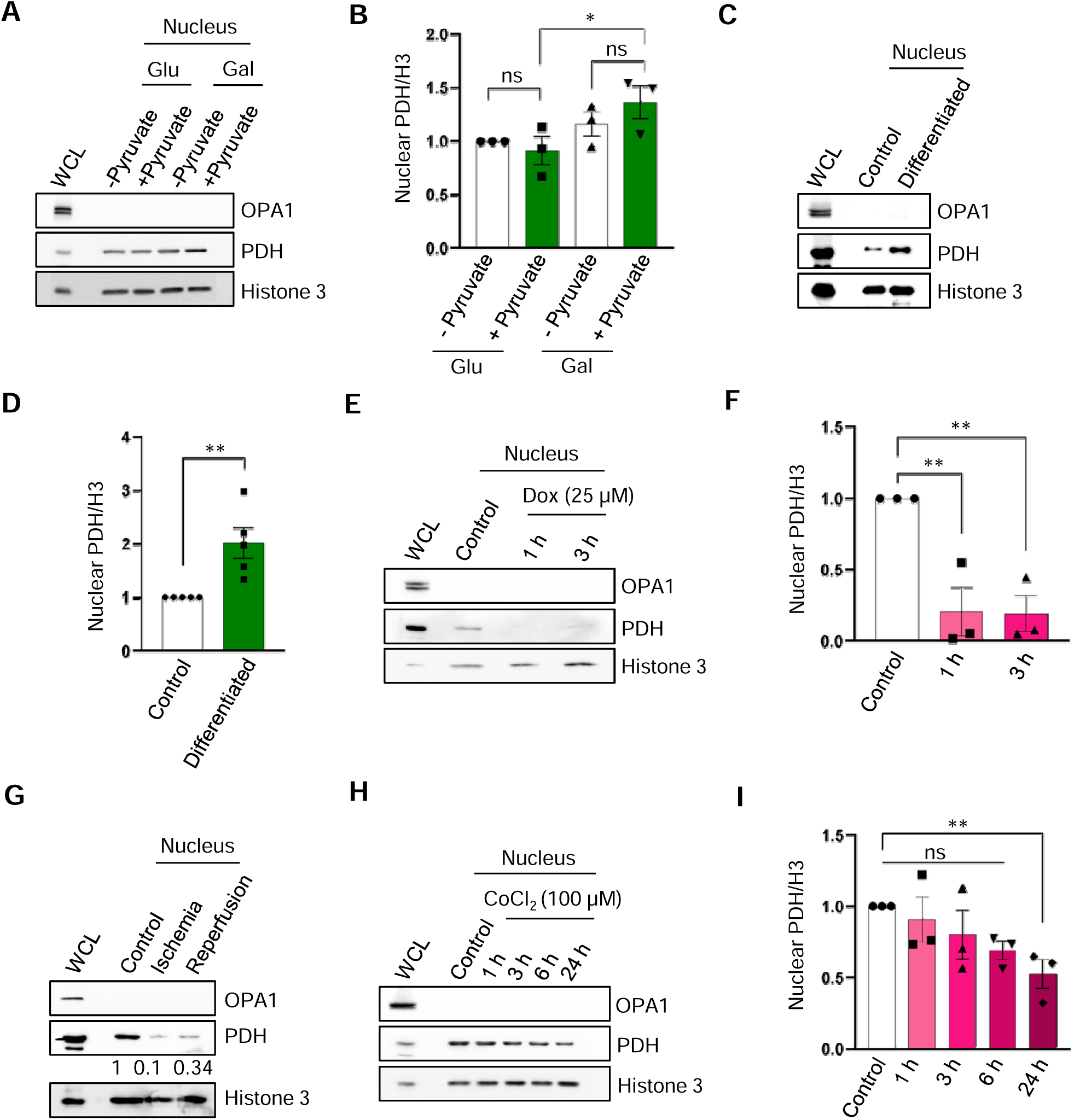
Regulation of nuclear PDH in basal and pathophysiological conditions. (A,B) H9C2 cardiomyoblasts were cultured in 25 mM glucose and 20 mM galactose-containing media (with a low 5 mM glucose) in the presence or absence of pyruvate (1 mM) for 24 hours; (A) Immunoblot of nuclear PDHE1, (B) Quantification of nuclear PDH (n=3). Nuclear PDH was normalized to Histone 3. WCL: Whole cell lysate, Glu: Glucose, Gal: Galactose. RM one-way ANOVA followed by Bonferroni post-hoc analysis, *p-value< 0.05, ns: non-significant. Error bars, SEM. (C,D) H9C2 cardiomyoblasts were differentiated into cardiomyocytes using retinoic acid (1 µM) for 5 days;(C) Immunoblot of nuclear PDHE1, (D) Quantification of nuclear PDH (n=5). Nuclear PDH was normalized to Histone 3, WCL: Whole cell lysate. Paired t-test, **p-value< 0.01. Error bars, SEM. (E,F) H9C2 cardiomyoblasts were treated with Doxorubicin (25 µM) for 1 hour and 3 hours. (E) Immunoblot of nuclear PDHE1, (F) Quantification of nuclear PDH (n=3), Nuclear PDH was normalized to Histone 3, WCL: Whole cell lysate. One-way ANOVA followed by Bonferroni post-hoc analysis, **p-value< 0.01. Error bars, SEM. (G) Immunoblot of nuclear PDHE1 in ischemia condition (2 hours normoxia followed by 2 hours hypoxia) and reperfusion condition (2 hours hypoxia followed by 2 hours normoxia). WCL: Whole cell lysate. (H,I) H9C2 cardiomyoblasts were treated with CoCl_2_ (100 µM) to mimic hypoxia conditions. Immunoblot of nuclear PDH E1-α. (I) Quantification of nuclear PDH (n=3). Nuclear PDH was normalized to Histone 3. Paired t-test, **p-value< 0.01, ns: non-significant. Error bars, SEM.

Further we were interested to see the status of PDH transit to the nucleus under differentiation of H9C2 cardiomyoblast to cardiomyocytes-like cells. Firstly, we confirmed that indeed differentiation is happening upon retinoic acid and 1% serum growth. Cells change in morphology and appear more flattened after differentiation as can be seen in phase contrast and phalloidin stained images (Figures S5C and S5D). Differentiation was also confirmed by an increase in the percentage of multinucleate cells (data not shown). Next, we checked the number of PDH^+^ MDVs in control and differentiated cells and observed an increase PDH^+^ MDVs in our system (Figures S5M and S5N) as has been observed previously [4]. Next, we queried whether nuclear PDH pool responds to this change in cell state. Indeed, we observed a twofold increase in nuclear PDH upon differentiation compared to control cells (Figures 5C and 5D). Previous reports showed that mitochondrial mass increases in the process of H9C2 differentiation [48]. Following the same trend, we also observed an increase in total cellular PDH levels (Figures S5E and S5F). Together our results suggest that nuclear PDH levels increase upon growth in galactose media and the differentiation process but not on pyruvate supplementation.

Several stress conditions that lead to mitochondrial depolarization increase MDV numbers. We next investigated the nuclear PDH levels under various stress conditions relevant to cardiac cells like doxorubicin treatment, ischemia-reperfusion injury and hypoxia. Doxorubicin, a cytotoxic drug, induces cardiotoxicity by causing mitochondrial stress and damaging the mitochondria [49]. The mitochondrial stress is initiated by production of superoxide, hydroxyl radicals, peroxynitrite etc [50]. We treated H9C2 cells with 25 µM doxorubicin [4] for 1 hour and 3 hours as significant cell death was observed only by 6 hours (Figure S5K). Nuclear PDH decreased drastically at 1 hour and 3 hours treatment of doxorubicin (Figures 5E and 5F). We confirmed that the total cellular PDH level remains unchanged (Figures S5G and S5H). Interestingly, PDH^+^ MDVs biogenesis also increased in doxorubicin treatment in both 1 and 3 hours (Figures S5O and S5P). This result shows that PDH^+^ MDVs is an early response to doxorubicin-induced stress but these MDVs are not targeting the nucleus to enable mito-nuclear PDH transit. Ischemia-reperfusion injury is one of the major causes of heart failure. Ischemia is caused by hypoxia and nutrient deficiency, followed by reperfusion through reoxygenation process. The reperfusion process produces free radicals, which damages the mitochondria [51]. We observed a reduction in nuclear PDH during ischemia and reperfusion compared to control conditions (Figure 5G). Our confocal data showed that the number of PDH^+^ MDVs has increased during reperfusion, but no significant increment observed in ischemia condition (Figures S5Q and S5R). Here again the generated MDVs do not target the nucleus but perhaps other cellular destinations with a reduction in PDH levels in the nucleus. Probably the PDH already present in the nucleus is turned over with no fresh entry through MDVs under these conditions.

Cobalt chloride is a chemical agent which helps to stabilize and accumulate HIF-1α to mimic hypoxia conditions [52]. Decreased nuclear PDH was observed in HeLa cells upon prolonged hypoxia [33]. To understand the effect of hypoxia on nuclear PDH levels in H9C2, we treated the cells with 100 µM cobalt chloride (CoCl_2_) to establish hypoxic conditions for different time intervals. We verified HIF1α stabilization under these conditions (data not shown). No cell death was observed even after 24 hours of CoCl_2_ treatment (Figure S5L). Nuclear transit of PDH decreased at 24 hours of CoCl_2_ treatment (Figures 5H and 5I), although total cellular PDH pool remained unchanged (Figures S5I and S5J).

Our findings suggest that PDH^+^ MDVs increase under most of the mitochondrial stress conditions as expected, but it is not targeted to the nucleus and the PDH levels in the nucleus remain unchanged as with experiments in Figure 3 for Antimycin A or reduce as shown here. However, the PDH^+^ MDVs generated under basal physiological conditions that necessitate a higher mitochondrial functionality like growth in galactose media or differentiation are targeted to the nucleus.

## Discussion

Our study investigates the retrograde communication pathway between mitochondria and nucleus through mitochondria-derived vesicles (MDVs). We show that PDHE1 is transported to the nucleus through MDVs. PDH is a large protein complex and we are currently investigating if other components of the complex like PDHE2 and E3 are also trafficked to the nucleus using MDVs to form a functional active complex. Our proteomics data identified multiple subunits of PDC complex as NEM sensitive MDV cargo proteins. Further we identified and validated COX4 as a novel MDV cargo protein.

Several mitochondrial proteins have moonlighting functions in the nucleus [24]. Multiple TCA cycle and ETC enzymes were found in the nucleus [25]. It is reported that PDH is present in the nucleus of different cell lines like A549, BaF3 and 3T3-L1 adipocyte cells [28, 42, 43]. We found PDH in the nucleus of rat heart and H9C2 cardiomyoblast lines whereas trace amount of PDH is also present in HEK293T cell nucleus as reported previously [29]. Moreover, we identified COX4 as present in the nucleus of HEK293T cells but not the cardiac cells suggesting cell type specificity in the process of mito-nuclear protein transit. The roles of trafficked proteins in helping mito-nuclear communication and homeostasis are also therefore likely to be cell type specific.

With our live cell imaging data, we show that PDH^+^ MDV-like structures that are tracker negative fuse with the nucleus but the fusion mechanism is still unknown. A previous study has shown that the entire PDH complex enters the nucleus through the lamina following MFN2 mediated direct mito-nuclear tethering [29]. However, in mature cardiac cells with spheroid, relatively less motile mitochondria embedded within the myofibrils [53], direct tethering might be difficult to achieve. We believe that in this context MDVs may mediate the transit of proteins like PDH. Using a reconstitution approach, we observe that nuclear PDH levels increase upon incubation with MDVs and this is sensitive to the presence of NEM, the inhibitor of NSF and vesicle fusion process. Interestingly the endogenous nuclear pools of NDUFS1 and VDAC do not change suggesting that their nuclear pools aren’t dependent on MDV mediated protein transit and some of these could be shuttled directly from the cytosol through the nuclear pore complex. Our reconstitution results in tandem with literature [28] suggests that while PDH requires mitochondrial entry to fold and form a functional complex following the cleavage of its mitochondrial localization signal; this enzyme further transits into the nucleus through MDVs to enable other moonlighting functions to take place. While we envisage trafficking to be the likely mode of PDH transit in our system our data does not rule out the possibility of protein transits across mito-nuclear tethers reported recently [29, 54].

Our results indicate that PDH^+^MDV biogenesis is actin dependent (Figure 3). We also show that like Tom20^+^ MDVs which were reported to be microtubule dependent [14], PDH+ MDV biogenesis also appears to be tubulin dependent. Actin mediates all endocytic budding events [55] and it would be interesting to further study how actin mediated force generation helps in MDV budding from the mitochondrial surface. Role of actin in MDV budding has not been explored. We need to decouple budding versus transit to understand the role of cytoskeletal elements in carrying MDVs to the nucleus.

Our preliminary observations suggest that global acetyl histone (H3K9/14Ac) levels in cardiac cells do not parallel the increase or decrease in nuclear PDH (data not shown). Unlike previous reports [27], this nuclear transit of PDH is also not stress driven (Figure 5) and happens more predominantly in basal states with higher mitochondrial oxidative phosphorylation. This is contrary to the observation on several mitochondrial dehydrogenases including PDHE1 translocating to the nucleus post chemotherapy-induced cardiotoxicity [27]. This study uses hiPSC-cardiomyocytes and further they are glucose deprived to get a pure cardiomyocytes line eliminating other cell types. In contrast, our studies are on shorter time points of mitochondrial stress in rat cardiomyoblasts.

We found that PDH^+^ MDV biogenesis increases under most of the mitochondrial stress conditions as reported previously [4], but this does not result in an increased nuclear pool of PDH. Surprisingly in most cases there is a reduction in nuclear pools commiserate with nuclear protein turnover once delivery of new cargo is stopped. Previous literature demonstrates syntaxin-17, a mitochondrial SNARE protein, associates with mature MDVs to mediate MDV-endolysosome fusion [9]. This indicates that appropriate v-SNARES that mediate MDV to nuclear membrane fusion have to be incorporated in the outer membrane of the subpopulation of PDH MDVs that deliver PDH to the nucleus. PDH itself, depending on cellular context, in Parkin dependent and independent manner localizes to different cellular compartments like the lysosome or multivesicular bodies (MVB) respectively [11]. This observation indicates that various destinations are plausible for PDH^+^ MDVs. We hypothesized that stress induced PDH packaging into MDVs is probably destined to the endolysosomal pathway and may be a quality control arm whereas basal trafficking to the nucleus could be to enable other moonlighting functions which require further probing.

Further, we establish a novel mito-nuclear trafficking route which could be employed by other mitochondrial proteins that have dual roles in both compartments but are functional only after mitochondrial import. Since MDVs form an ancient vesicle compartment [19] we envisage MDV mediated nuclear trafficking of proteins as a general means of the endosymbiont communicating and controlling the host nuclear program.

## Key Resource Table

| REAGENT or RESOURCE | SOURCE | IDENTIFIER |
| --- | --- | --- |
| Antibodies |  |  |
| ACP/NDUFAB1 | Cell Signaling Technology | Cat#71814, RRID: AB_3086728 |
| COX1 | Cell Signaling Technology (CST) | Cat# 55159T |
| COX10 | Cell Signaling Technology (CST) | Cat# 24744T |
| COX4 (3E11) Rabbit mAb | Cell Signaling Technology (CST) | Cat# 4850, RRID: AB_2085424 |
| COX4 Polyclonal Antibody | Thermo Fisher Scientific Invitrogen | Cat# PA5-19471, RRID: AB_11004909 |
| Cytochrome c (136F3) Rabbit mAb | Cell Signaling Technology (CST) | Cat# 4280, RRID: AB_10695410 |
| Histone 3 | Cell Signaling Technology (CST) | Cat# 9715 |
| Histone 3 | Santa Cruz | Cat# 517576 |
| HSP60 (D6F1) XP® Rabbit mAb | Cell Signaling Technology (CST) | Cat# 12165, RRID: AB_2636980 |
| OPA1 Mouse mAb | BD Biosciences | Cat# 612606, RRID: AB_399888 |
| PHB1 Polyclonal Antibody | Cell Signaling Technology (CST) | Cat# 2426, RRID: AB_823689 |
| Pyruvate Dehydrogenase (C54G1) Rabbit mAb | Cell Signaling Technology (CST) | Cat# 3205, RRID: AB_2162926 |
| SDHA (D6J9M) Rabbit mAb | Cell Signaling Technology (CST) | Cat# 11998, RRID: AB_2750900 |
| SOD1 (71G8) Mouse mAb | Cell Signaling Technology (CST) | Cat# 4266, RRID: AB_2193898 |
| TOM20 (D8T4N) Rabbit mAb | Cell Signaling Technology (CST) | Cat#42406, RRID: AB_2799825 |
| Tubulin | Merck | Cat#AB9354 |
| UQCRFS1/RISP | Cell Signaling Technology (CST) | Cat# 95231, RRID: AB_2922784 |
| VDAC (D73D12) Rabbit mAb | Cell Signaling Technology (CST) | Cat#4661, RRID: AB_10557420 |
| Chemicals, peptides, and recombinant proteins |  |  |
| Acetone | Himedia | MB179 |
| Acrylamide | BIO RAD | 161-0101 |
| 2 mercaptoethanol | Himedia | MB041 |
| ADP disodium salt | Himedia | MB191 |
| Ammonium persulphate | Himedia | MB003 |
| Antibiotic Antimycotic solution | Sigma Aldrich | A5955 |
| Antimycin A | Sigma | A8674 |
| ATP disodium salt | Himedia | RM439 |
| Bis-acrylamide | Himedia | MB005 |
| Bromophenol blue | Himedia | MB123 |
| Cobalt chloride | Merck | S4321140 |
| D- Glucose | Sigma Aldrich | G8270 |
| DAPI | Himedia | MB097 |
| D-Galactose | Sigma Aldrich | G0750 |
| Dipotassium hydrogen phosphate | Himedia | TC596 |
| di-Sodium hydrogen phosphate dodecahydrate | Himedia | GRM1418 |
| DMEM low glucose Medium | Himedia | AL006A |
| DMEM Medium | Sigma Aldrich | D0547 |
| DMSO | Sigma Aldrich | D2438 |
| Dodecyl sulphate sodium salt | Himedia | TC698 |
| Doxorubicin | Sigma Aldrich | 44583 |
| EGTA | Himedia | GRM1530 |
| EM grade glutaraldehyde | Merck | G5882 |
| Endothelial cell growth medium | Sigma Aldrich | 211-500 |
| Fetal Bovine Serum | Gibco | 10270106 |
| Fluoromount G | Invitrogen | 4958-02 |
| Formvar/Carbon 200 Mesh, Gold grid | Electron Microscopy | FCF200-AU25 |
|  | Sciences |  |
| Glycerol | Himedia | TC503 |
| Glycine | Himedia | MB013 |
| HEPES | Himedia | MB016 |
| Hydrocortisone | Sigma Aldrich | H0888 |
| Insulin | Sigma Aldrich | 19278 |
| Latrunculin B | Sigma Aldrich | 428020 |
| Lipofectamine 2000 | Invitrogen | 11668030 |
| M-199 Medium | Sigma Aldrich | M0393 |
| Magnesium acetate | Himedia | MB039 |
| Mannitol | Himedia | MB198 |
| Mdivi | Sigma Aldrich | M1099 |
| Methanol | Thermo Fisher<br>Qualigens | Q43637G |
| MitoTracker | Cell Signaling<br>Technology (CST) | 8778S |
| N-Ethylmaleimide | Sigma | E3876 |
| Nocodazole | Sigma Aldrich | M1404 |
| NP-40 solution | Himedia | MB143 |
| Opti-MEM Medium | Gibco | 31985070 |
| Phosphotungstic acid hydrate | Sigma Aldrich | 79690 |
| Potassium chloride | Himedia | GRM3934 |
| Potassium dihydrogen orthophosphate monobasic | Himedia | GRM3943 |
| Retinoic acid | Sigma Aldrich | R2625 |
| Sodium chloride | Himedia | GRM3954 |
| Sodium pyruvate | Sigma Aldrich | S8636 |
| Sodium succinate | Himedia | GRM418 |
| Sucrose | Himedia, Sigma | PCT0616, S0389 |
| TEMED | BIO RAD | 161-0800 |
| Tris-Base | BIO RAD, Himedia | 1610719, TC072 |
| Trypsin-EDTA solution | Sigma Aldrich | T4174 |
| Tween-20 | Himedia | TC287 |
| Recombinant DNA |  |  |
| pLV[Exp]-Puro-CMV>hPDHA1[NM_001173454.2]/mCherry | Vector Builder | VB221211-1356azz |
| Experimental models: Organisms/strains |  |  |
| Rat (Wistar) |  |  |
| Software and algorithms |  |  |
| Fiji | NIH | https://fiji.sc/ |
| GraphPad Prism | GraphPad Software<br>Inc | http://www.graphpad.com/ |

## Materials and Methods

### Cell culture and treatments

Rat embryonic cardiomyoblasts (H9C2), Human kidney cells (HEK293T) (ATCC) and Human umbilical vein endothelial cells (HUVEC) (Promocell) were used in this study. H9C2 cardiomyocytes and HEK293T cells were maintained in Dulbecco’s modified eagle medium (DMEM) (Sigma-Aldrich) supplemented with 10% FBS (Gibco) and 1% antibiotic antimycotic solution (Sigma-Aldrich). The H9C2 cardiomyoblasts were additionally supplemented with 1 mM sodium pyruvate except in experiments where the role of pyruvate is tested (Sigma-Aldrich). HUVEC was maintained in endothelial cell growth media (Sigma-Aldrich) supplemented with 2% FBS and 2% antibiotic antimycotic solution (Sigma-Aldrich).

All cell culture chemicals are from Sigma Aldrich unless otherwise indicated. Please find details in the key resource table. Cell treatments are indicated in the respective figures, as appropriate, and were: antimycin A (20 μM, Sigma-Aldrich, 1h), latrunculin B (1 μM, 15 min), nocodazole (10 μM, 30 min), doxorubicin (25 μM, 1h, 3h), cobalt chloride (100 µM, Merck, 1h-24h). Appropriate time matched controls were set for every treatment condition. To investigate the effect of galactose, H9C2 cardiomyoblasts were cultured in DMEM low glucose (5 mM glucose, Himedia) with galactose (20 mM) for 24 hours.

### Lentiviral transduction

To prepare stable cell line overexpressing PDH mCherry, initially viral particles were produced in HEK293T. HEK293T was transfected with pLVX PDH mCherry and packaging vectors, pMDG2 and pAX2 using lipofectamine 2000. After 48 hours, viral supernatant was added to H9C2 for 4 hours. Cells were selected for stable expression using puromycin (1.5 _μ_g/ml).

### H9C2 cardiomyoblast differentiation

H9C2 cardiomyoblasts were differentiated using retinoic acid in low serum containing media [56]. All-trans retinoic acid (Sigma-Aldrich) was prepared in DMSO and stored in dark at - 20°C. A total of 81,000 cells for non-differentiation and 3,00,000 cells for differentiation were plated in 60 mm tissue culture dish and cultured one day in 10% FBS containing media in order to adhere cells. Non-differentiated cell plate was maintained in 10% FBS containing media. For differentiation, cells were cultured with 1µM all-trans retinoic acid in low serum containing media (1% FBS) for 5 days. On the 6th day, cells were harvested for nuclei isolation or whole cell lysate preparation following described procedures.

### Ischemia-Reperfusion induction

H9C2 cardiomyoblasts were seeded in 100 mm tissue culture dishes. For ischemia conditions, cells were incubated with complete media in a 5% CO2 incubator for 2 hours, followed by incubation in hypoxia chamber (New Brunswick Galaxy 48R) with glucose free and serum free media for 2 hours. To establish reperfusion conditions, cells were incubated in hypoxia conditions as mentioned above, followed by incubation in a regular 5% CO_2_ incubator with complete media for 2 hours. Post-treatment, cells were subjected to nucleus isolation following the described procedures.

### Isolation of MDVs

The protocol followed as well as appropriate controls is detailed in Nair et al., 2024 [40]. Cardiac MDVs were budded from mitochondria isolated from rat hearts.

The budding reaction solution was constituted of 1 mM ATP, 5 mM succinate, 80 μM ADP and 2 mM K_2_HPO_4_, in 1X MIB. A parallel reaction containing NEM (20 mM) was also set up. Budding reactions were incubated at 37°C for 2 hours in the incubator with shaking. The budding reactions were centrifuged at 10,000g for 10 minutes and the supernatant containing the MDVs was collected.

To further purify the vesicles, MDVs were floated by sucrose gradient centrifugation (Beckman Coulter - Sw41Ti rotor). Samples were loaded at the bottom of the discontinuous sucrose gradient, by loading 0.5 ml of sample to 1.5 ml of 80% sucrose, resulting in 60% sucrose. The gradient from the bottom was as follows: 60%, 50%, 40%, 30%, 20%, and 10 % (2ml each). Samples were centrifuged at 32,500 rpm (100,000g) for 16 hours at 4°C and 1 ml of the fractions were collected from the top.

### Proteomics analysis

All budding reactions were concentrated using 3kDa Amicon filter column to 50 μL and the concentrated samples were loaded on SDS PAGE. The samples were cut from gel upon reaching the interface between the separating gel and the resolving gel. The gel piece was destained and 100% acetonitrile solution was used to dry the gel. This was followed by overnight digestion with 13 ng/μL trypsin in 50 mM ammonium bicarbonate. 5% formic acid/acetonitrile (1:2, v/v) extraction buffer was used to extract peptide from the gel. Finally, the samples were reconstituted in 3% acetonitrile.

A nano ACQUITY UPLC System (Waters) chromatography system was used for the LC analysis for the first two sets of samples. The Mass Lynx 4.1 SCN781 software was used for the acquisition. The peptides were separated by reversed-phase chromatography. MS analysis of eluting peptides was carried out on a SYNAPT® G2 High-Definition MS™ system (HDMS^E^ System) (Waters). The analyses were carried out in positive mode electrospray ionization (ESI). The third set of samples were analysed using Orbitrap Eclipse. Xcalibur software was used for data acquisition. Protein identification was done by analysing LC-MS data using Progenesis QI for Proteomics V4.2 (Waters) for the initial two MS runs and Proteome Discoverer v2.5 was used for the last MS run. All three runs were separated on a binary phase system: the mobile phase-A was water, and the mobile phase-B was acetonitrile; both contain 0.1% formic acid. The proteins were identified from the *Rattus norvegicus* database on UniProt.

For further analysis, only proteins with at least one unique peptide were considered. Mitochondrial localization of the protein was established with the help of MitoCarta 3.0 and The Integrated Mitochondrial Protein Index (IMPI) database. The sublocalization of these proteins within the mitochondria was established with the help of these databases. In addition to that a literature survey done by us gave evidence for the mitochondrial sublocalization. To identify the high confidence NEM sensitive MDV cargo, we compared the control proteins with the NEM proteins, and the proteins detected only in the control were considered. A Reactome pathway analysis of the confirmed MDV cargo was performed. Our MDV proteome was compared to the Reactome database including the interaction data from IntAct.

### TEM imaging

The budding reactions were loaded onto gold Formvar carbon coated TEM grid (FCF200-Au; Electron Microscopy Sciences) and incubated overnight at 4°C. To fix, 2.5% glutaraldehyde was added for 30s and negative staining was done using uranyl acetate for 1 min. Images were acquired using JEM-1011 electron microscope (JEOL) using Digital Micrograph software.

### Isolation of Nucleus from Cells

Cells were grown on a 100 mm tissue culture dish for nucleus isolation. Cells were washed with ice-cold 1X PBS and scrapped in 0.1% NP-40 solution by the help of cell scrapper lysing only the plasma membrane. The sample was washed 2 times with 0.1% NP-40 solution by centrifugation at 700g at 4°C [57]. Then samples were loaded on top of the 60% sucrose cushion and processed for ultracentrifugation (Beckman Coulter - Sw41Ti rotor) at 30,000g for 1 hour at 4°C. After ultracentrifugation, the nuclear pellet was resuspended in Laemmli sample buffer and ultrasonicated to shear the genomic DNA before loading for western blots.

### Isolation of nucleus from rat heart

Nucleus isolation from rat heart was adapted from previously established protocols [57, 58]. Rat hearts were collected in ice-cold PBS. The hearts were chopped and followed by centrifugation at 3000g for 10 min at 4°C. Supernatant was discarded and the pellet was washed in PBS by centrifugation. The pellet was resuspended in isolation buffer (Mannitol 220 mM sucrose 68 mM; KCl 80 mM; EGTA 0.5 mM; Mg(C□H□O□)□2 mM; HEPES, pH 7.4 20 mM) and homogenized using a 50 ml Dounce tissue homogenizer, followed by centrifugation at 3000g for 10 min at 4°C. The pellet was resuspended in isolation buffer containing 0.1% NP-40 and passed through a 40 μm cell strainer (Himedia). The filtered suspension was washed by centrifugation at 500g for 10 min at 4°C. The resuspended pellet was loaded on a 60% sucrose cushion. This was followed by ultracentrifugation using swinging bucket rotor (Beckman Coulter - Sw41Ti rotor) at 30,000g for 1 hour at 4°C. The nuclear pellet was resuspended in 200 μL of 1X Laemmli buffer for western blotting or 200 μL of MIB for reconstitution experiment. Whole tissue lysate was prepared from rat heart tissue using 1X Laemmli buffer followed by sonication.

### Immunoblotting

All the samples were prepared in 1X Laemmli buffer, sonicated and boiled at 95°C for 5 min. Following separation of proteins by SDS-PAGE, the proteins were transferred to nitrocellulose membrane. Ponceau S staining was done to determine efficient transfer and equal load. The blots were immunoblotted with respective primary and secondary antibodies. The primary antibodies used were anti-VDAC (1:1000) (CST 4661T), anti-TOM20 (1:1000) (CST 42406S), anti-OPA1 (1:2000) (BD Biosciences cat 612606), anti-PDH (1:1000) (CST 3205S), anti-SDHA (1:1000) (CST 11998T), anti-COX4 (1:1000) (CST 3E11 4850S), anti-COX4 (1:2000) (Invitrogen PA5-19471), anti-PHB1 (1:1000) (CST 2426T), anti-cytochrome c (1:1000) (CST 4280t), anti HSP60 (1:1000) (CST), anti SOD1 (1:1000) (CST 4266T), anti-histone 3 (1: 5000) (Santa Cruz sc517576), anti-histone 3 (1:5000) (CST 9715), anti- -β tubulin (1:2000) (Merck), anti-NDUFS1 (1:1000) (CST 70264T), anti-COX10 (1:1000) (CST 24744T), anti-COX1 (1:1000) (CST 55159T), anti-ACP/NDUFAB1 (1:1000) (CST 71814T), anti-UQCRFS1/RISP (1:1000) (CST 95231T). The secondary antibodies are Goat anti-rabbit (1:10,000) (CST) and Goat anti-mouse (1:10,000) (CST). The protein bands were detected using ECL solution and developed either X ray or on a Chemidoc machine (ImageQuant LAS500)

### Antimycin A washout assay

70,000 H9C2 cardiomyoblasts were seeded on coverslips in 35 mm dishes. All the cells were pre-treated with Drp1 inhibitor, Mdivi (Sigma-Aldrich) (1μM) for 1 hour and Mdivi is present throughout the experiment. The control cells were treated with DMSO for 1 h while the treatment cells were treated with antimycin A (20 μM) for 1 h. After treatment the antimycin containing media was removed and Mdivi containing media was added. The treated cells were fixed at 0 min, 30 min, 60 min, 90 min and 120 min after antimycin washout. The cells were fixed with ice-cold methanol-acetone (1:1) for 5 min at −20°C.

### Immunofluorescence

Cells were grown on coverslip and treated according to each experiment. Post-treatment 500nM MitoTracker (CST) was added to stain mitochondria and incubated at 37°C for 30 minutes. Cells were washed 3 times with 1X PBS. In order to fix the cells, ice-cold 100% methanol was added and incubated in −20°C for 15 minutes. Then cells were washed 3 times with 1X PBS and blocked with 1% BSA solution at room temperature for 1 hour. Cells were incubated with primary antibodies for 1 hour at room temperature and subsequent 3 washes with 1X PBS. The primary antibodies used were anti-PDH E1-α (1:25) (Santacruz, sc-377092), anti-COX4 (1:100) (CST 3E11 4850S) and anti-tubulin (1:200) (). Then, cells were incubated with respective Alexa Fluor secondary antibodies (Jackson Immunoresearch) for 1 hour at room temperature in dark, followed by 3 washes with 1X PBS. Actin was stained with phalloidin (1:400) (CST). Cells were counterstained with DAPI (Himedia) (1:10, 000 dilution) for 5 minutes at room temperature, followed by 5 times washing with 1X PBS. Finally, coverslips were mounted with Fluoromount G (Invitrogen).

### Imaging

Confocal images (∼0.3 μm step size) were acquired with Olympus Fluoview FV3000 60X, Oil. The excitation wavelengths used were 405, 488, 561 and 640 nm. Image files were analysed using ImageJ.

For live imaging, H9C2 cells overexpressing PDH-mCherry were seeded on a 35mm glass bottom dish. Cells were tracked with MitoTracker Far Red (500nM) for 30 min prior to imaging followed by three PBS washes. The nucleus was stained with Hoechst followed by five PBS washes. Cells were imaged at 37°C in a chamber with 5% CO_2_. Images were acquired with Olympus Fluoview FV3000.

### *In vitro* reconstitution

To study cargo delivery into nucleus by MDVs, we established cell free reconstitution reactions between nucleus and MDVs. Nucleus and cardiac MDVs were isolated in parallel from rat heart as described above. We reconstituted the nucleus and MDVs in the presence and absence of NEM (20 mM). A nucleus only reaction was set up as a control. All the reactions occurred in the presence of 0.5 μM GTP (Sigma-Aldrich). The reactions were incubated at 37°C for 1h in a shaking water bath. The nucleus was re-isolated by ultracentrifugation and the samples were prepared in 1X Laemmli buffer.

### Quantification and Statistical Analysis

For western blots, band intensities were quantified using ImageJ software. All statistical analyses were performed using GraphPad Prism software (version 8.4.3). Student t-test was performed to compare fold changes against the normalized control to compare two samples. Ordinary one-way ANOVA or repeated measure one-way ANOVA followed by Bonferroni post-hoc analysis was performed to analyze a single independent variable with three or more samples. Error bar is reported as mean ± SEM. ****p-value< 0.0001, ***p-value< 0.001, **p-value< 0.01, *p-value< 0.0 5.

## Ethics statement

All animal experiments were performed after valid ethical clearance from the Institutional (RGCB) Animal Bioethics Committee (IAEC/867/ANANTHA/2021 and IAEC/940/ANANTHA/2023) which is registered under Committee for the Purpose of Control and Supervision of Experiments on Animals (CPCSEA), Govt. of India [326/GO/ReBiBt/S/2001/CPCSEA].

## Data Availability

All relevant data supporting the key findings of this study are availa ble within the article and the supplementary data file S1 and S2. All data reported in this paper will be shared by lead contact/ corresponding author (AS) upon request.

## Acknowledgements

This work is supported by the Ramalingaswami Re-entry Fellowship (BT/RLF/Re-entry/51/2019 and SERB POWER grant (SPG/2021/003267) awarded to AS by the Department of Biotechnology (DBT) and Science and Engineering Research Board, Department of Science and Technology (DST), Government of India respectively, and JRF fellowship from University Grants Commission to TR. Authors are grateful to the confocal, proteomics and animal research facilities as well as the Instrumentation Department at Rajiv Gandhi Centre for Biotechnology (RGCB) for their help with experiments.

## Author contributions

Conceptualization, A.S. and T.R.

Methodology T.R., D.S., S.B., L.R., V.S.I, N.N, A.M., M.J., S.G., J.B.J., A.S.

Investigation T.R., D.S., S.B., L.R., V.S.I, N.N, A.M., M.J., S.G., J.B.J., A.S.

Writing – Original Draft, A.S, T.R, D.S., Writing – Review & Editing, all authors Funding Acquisition A.S.

Supervision A.S.

## References

1. Thapa, D., et al., Adropin regulates pyruvate dehydrogenase in cardiac cells via a novel GPCR-MAPK-PDK4 signaling pathway. Redox Biol, 2018. 18: p. 25–32.

2. Cadete, V.J.J., et al., Mitochondrial quality control in the cardiac system: An integrative view. Biochim Biophys Acta Mol Basis Dis, 2019. 1865(4): p. 782–796.

3. Soubannier, V., et al., A vesicular transport pathway shuttles cargo from mitochondria to lysosomes. Curr Biol, 2012. 22(2): p. 135–41.

4. Cadete, V.J., et al., Formation of mitochondrial-derived vesicles is an active and physiologically relevant mitochondrial quality control process in the cardiac system. J Physiol, 2016. 594(18): p. 5343–62.

5. Dorn, G.W., 2nd, *Mitochondrial dynamism and heart disease: changing shape and shaping change*. EMBO Mol Med, 2015. 7(7): p. 865–77.

6. Neuspiel, M., et al., Cargo-selected transport from the mitochondria to peroxisomes is mediated by vesicular carriers. Curr Biol, 2008. 18(2): p. 102–8.

7. Braschi, E., et al., Vps35 mediates vesicle transport between the mitochondria and peroxisomes. Curr Biol, 2010. 20(14): p. 1310–5.

8. McLelland, G.L., et al., Parkin and PINK1 function in a vesicular trafficking pathway regulating mitochondrial quality control. Embo j, 2014. 33(4): p. 282–95.

9. McLelland, G.L., et al., Syntaxin-17 delivers PINK1/parkin-dependent mitochondrial vesicles to the endolysosomal system. J Cell Biol, 2016. 214(3): p. 275–91.

10. Matheoud, D., et al., Parkinson’s Disease-Related Proteins PINK1 and Parkin Repress Mitochondrial Antigen Presentation. Cell, 2016. 166(2): p. 314–327.

11. Todkar, K., et al., Selective packaging of mitochondrial proteins into extracellular vesicles prevents the release of mitochondrial DAMPs. Nat Commun, 2021. 12(1): p. 1971.

12. Ryan, T.A., et al., Tollip coordinates Parkin-dependent trafficking of mitochondrial-derived vesicles. Embo j, 2020. 39(11): p. e102539.

13. Juhász, G., A mitochondrial-derived vesicle HOPS to endolysosomes using Syntaxin-17. J Cell Biol, 2016. 214(3): p. 241–3.

14. König, T., et al., MIROs and DRP1 drive mitochondrial-derived vesicle biogenesis and promote quality control. Nat Cell Biol, 2021. 23(12): p. 1271–1286.

15. Abuaita, B.H., T.L. Schultz, and M.X. O’Riordan, Mitochondria-Derived Vesicles Deliver Antimicrobial Reactive Oxygen Species to Control Phagosome-Localized Staphylococcus aureus. Cell Host Microbe, 2018. 24(5): p. 625–636.e5.

16. Hazan Ben-Menachem, R., et al., Mitochondrial-derived vesicles retain membrane potential and contain a functional ATP synthase. EMBO Rep, 2023. 24(5): p. 17.

17. Andrade-Navarro, M.A., L. Sanchez-Pulido, and H.M. McBride, Mitochondrial vesicles: an ancient process providing new links to peroxisomes. Curr Opin Cell Biol, 2009. 21(4): p. 560–7.

18. Gould, S.B., S.G. Garg, and W.F. Martin, Bacterial Vesicle Secretion and the Evolutionary Origin of the Eukaryotic Endomembrane System. Trends Microbiol, 2016. 24(7): p. 525–534.

19. König, T. and H.M. McBride, Mitochondrial-derived vesicles in metabolism, disease, and aging. Cell Metab, 2024. 36(1): p. 21–35.

20. Lo, S.C. and M. Hannink, PGAM5 tethers a ternary complex containing Keap1 and Nrf2 to mitochondria. Exp Cell Res, 2008. 314(8): p. 1789–803.

21. Monaghan, R.M., et al., A nuclear role for the respiratory enzyme CLK-1 in regulating mitochondrial stress responses and longevity. Nat Cell Biol, 2015. 17(6): p. 782–92.

22. Yogev, O., et al., Fumarase: a mitochondrial metabolic enzyme and a cytosolic/nuclear component of the DNA damage response. PLoS Biol, 2010. 8(3): p. e1000328.

23. Gibellini, L., et al., Evidence for mitochondrial Lonp1 expression in the nucleus. Sci Rep, 2022. 12(1): p. 10877.

24. Monaghan, R.M. and A.J. Whitmarsh, Mitochondrial Proteins Moonlighting in the Nucleus. Trends Biochem Sci, 2015. 40(12): p. 728–735.

25. Nagaraj, R., et al., Nuclear Localization of Mitochondrial TCA Cycle Enzymes as a Critical Step in Mammalian Zygotic Genome Activation. Cell, 2017. 168(1-2): p. 210–223.e11.

26. Li, W., et al., Nuclear localization of mitochondrial TCA cycle enzymes modulates pluripotency via histone acetylation. Nat Commun, 2022. 13(1): p. 7414.

27. Srivastava, S., et al., Nuclear translocation of mitochondrial dehydrogenases as an adaptive cardioprotective mechanism. Nat Commun, 2023. 14(1): p. 023–40084.

28. Sutendra, G., et al., A nuclear pyruvate dehydrogenase complex is important for the generation of acetyl-CoA and histone acetylation. Cell, 2014. 158(1): p. 84–97.

29. Zervopoulos, S.D., et al., MFN2-driven mitochondria-to-nucleus tethering allows a non-canonical nuclear entry pathway of the mitochondrial pyruvate dehydrogenase complex. Mol Cell, 2022. 82(5): p. 1066–1077.e7.

30. Soubannier, V., et al., Reconstitution of mitochondria derived vesicle formation demonstrates selective enrichment of oxidized cargo. PLoS One, 2012. 7(12): p. e52830.

31. Lee, J., et al., The plasticity of the pyruvate dehydrogenase complex confers a labile structure that is associated with its catalytic activity. PLoS One, 2020. 15(12): p. e0243489.

32. Patel, M.S., et al., The pyruvate dehydrogenase complexes: structure-based function and regulation. J Biol Chem, 2014. 289(24): p. 16615–23.

33. Eguchi, K. and K. Nakayama, Prolonged hypoxia decreases nuclear pyruvate dehydrogenase complex and regulates the gene expression. Biochem Biophys Res Commun, 2019. 520(1): p. 128–135.

34. Chen, J., et al., Compartmentalized activities of the pyruvate dehydrogenase complex sustain lipogenesis in prostate cancer. Nat Genet, 2018. 50(2): p. 219–228.

35. Mocholi, E., et al., Pyruvate metabolism controls chromatin remodeling during CD4(+) T cell activation. Cell Rep, 2023. 42(6): p. 112583.

36. Vasam, G., et al., Proteomics characterization of mitochondrial-derived vesicles under oxidative stress. Faseb j, 2021. 35(4): p. e21278.

37. Towers, C.G., et al., Mitochondrial-derived vesicles compensate for loss of LC3-mediated mitophagy. Dev Cell, 2021. 56(14): p. 2029–2042.e5.

38. Roberts, R.F., et al., Proteomic Profiling of Mitochondrial-Derived Vesicles in Brain Reveals Enrichment of Respiratory Complex Sub-assemblies and Small TIM Chaperones. J Proteome Res, 2021. 20(1): p. 506–517.

39. Weidman, P.J., et al., Binding of an N-ethylmaleimide-sensitive fusion protein to Golgi membranes requires both a soluble protein(s) and an integral membrane receptor. J Cell Biol, 1989. 108(5): p. 1589–96.

40. Nidhi, N., et al., In Vitro Budding and Floatation based Enrichment of Mitochondria-derived Vesicles for Proteomics from Rat Heart. bioRxiv, 2024: p. 2024.03.04.583262.

41. Wang, S., et al., Prohibitin co-localizes with Rb in the nucleus and recruits N-CoR and HDAC1 for transcriptional repression. Oncogene, 2002. 21(55): p. 8388–96.

42. Chueh, F.Y., et al., Nuclear localization of pyruvate dehydrogenase complex-E2 (PDC-E2), a mitochondrial enzyme, and its role in signal transducer and activator of transcription 5 (STAT5)-dependent gene transcription. Cell Signal, 2011. 23(7): p. 1170–8.

43. Richard, A.J., H. Hang, and J.M. Stephens, Pyruvate dehydrogenase complex (PDC) subunits moonlight as interaction partners of phosphorylated STAT5 in adipocytes and adipose tissue. J Biol Chem, 2017. 292(48): p. 19733–19742.

44. Wu, X.S., et al., Actin Is Crucial for All Kinetically Distinguishable Forms of Endocytosis at Synapses. Neuron, 2016. 92(5): p. 1020–1035.

45. Schuh, M., An actin-dependent mechanism for long-range vesicle transport. Nat Cell Biol, 2011. 13(12): p. 1431–6.

46. Aguer, C., et al., Galactose enhances oxidative metabolism and reveals mitochondrial dysfunction in human primary muscle cells. PLoS One, 2011. 6(12): p. e28536.

47. Kase, E.T., et al., Remodeling of oxidative energy metabolism by galactose improves glucose handling and metabolic switching in human skeletal muscle cells. PLoS One, 2013. 8(4): p. e59972.

48. Comelli, M., et al., Cardiac differentiation promotes mitochondria development and ameliorates oxidative capacity in H9c2 cardiomyoblasts. Mitochondrion, 2011. 11(2): p. 315–26.

49. Osataphan, N., et al., Effects of doxorubicin-induced cardiotoxicity on cardiac mitochondrial dynamics and mitochondrial function: Insights for future interventions. J Cell Mol Med, 2020. 24(12): p. 6534–6557.

50. Wu, L., et al., Mitochondrial quality control mechanisms as therapeutic targets in doxorubicin-induced cardiotoxicity. Trends Pharmacol Sci, 2023. 44(1): p. 34–49.

51. Pedriali, G., et al., Perspectives on mitochondrial relevance in cardiac ischemia/reperfusion injury. Front Cell Dev Biol, 2022. 10: p. 1082095.

52. Prajapat, S.K., K.C. Maharana, and S. Singh, Mitochondrial dysfunction in the pathogenesis of endothelial dysfunction. Mol Cell Biochem, 2023.

53. Tepp, K., et al., High efficiency of energy flux controls within mitochondrial interactosome in cardiac intracellular energetic units. Biochim Biophys Acta, 2011. 1807(12): p. 1549–61.

54. Desai, R., et al., Mitochondria form contact sites with the nucleus to couple prosurvival retrograde response. Sci Adv, 2020. 6(51).

55. Galletta, B.J. and J.A. Cooper, Actin and endocytosis: mechanisms and phylogeny. Curr Opin Cell Biol, 2009. 21(1): p. 20–7.

56. Branco, A.F., et al., Gene Expression Profiling of H9c2 Myoblast Differentiation towards a Cardiac-Like Phenotype. PLoS One, 2015. 10(6): p. e0129303.

57. Nabbi, A. and K. Riabowol, Rapid Isolation of Nuclei from Cells In Vitro. Cold Spring Harb Protoc, 2015. 2015(8): p. 769–72.

58. Bergmann, O. and S. Jovinge, Isolation of cardiomyocyte nuclei from post-mortem tissue. J Vis Exp, 2012(65).

