## Supplementary figures for "Mitochondria-derived Vesicles mediate Mito-nuclear Trafficking of Cargo"

### Supplementary Figure 1

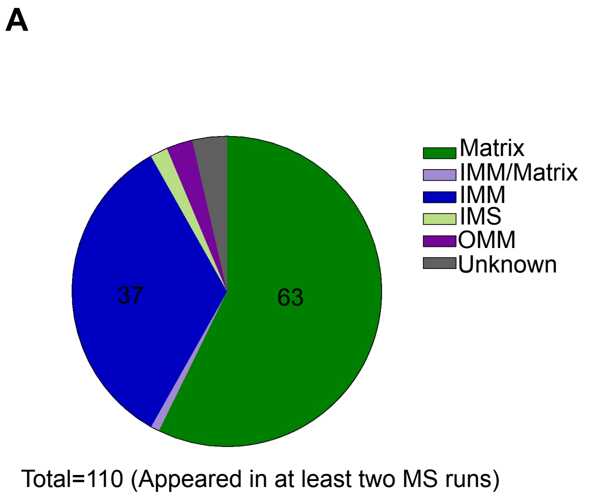


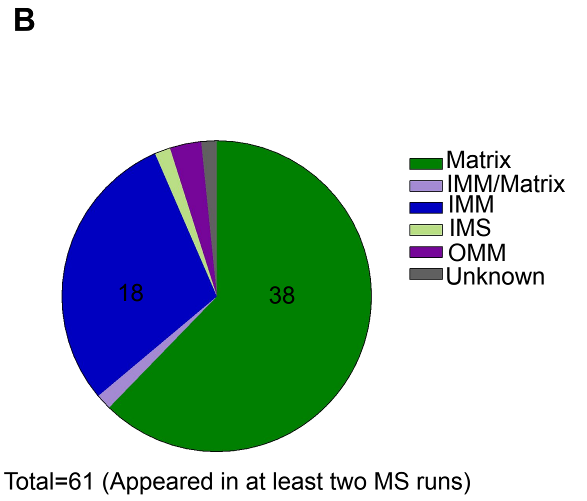


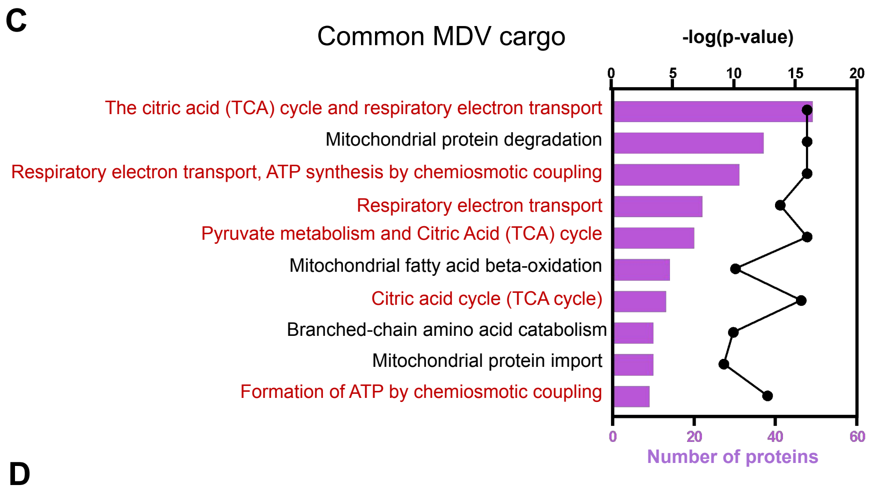


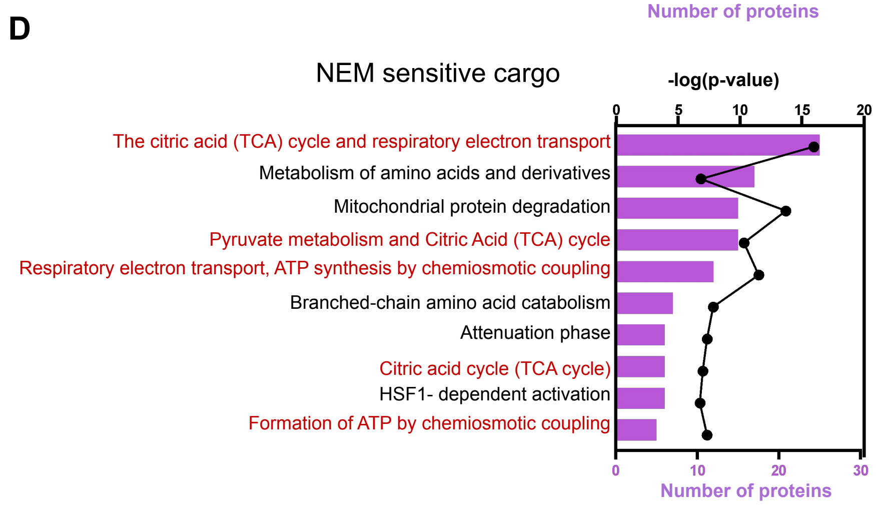


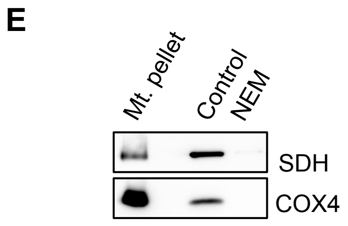


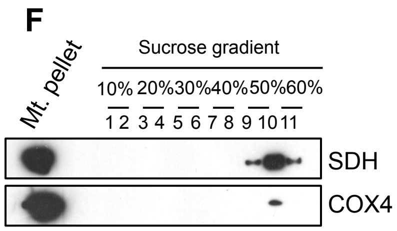

### Supplementary Figure 1

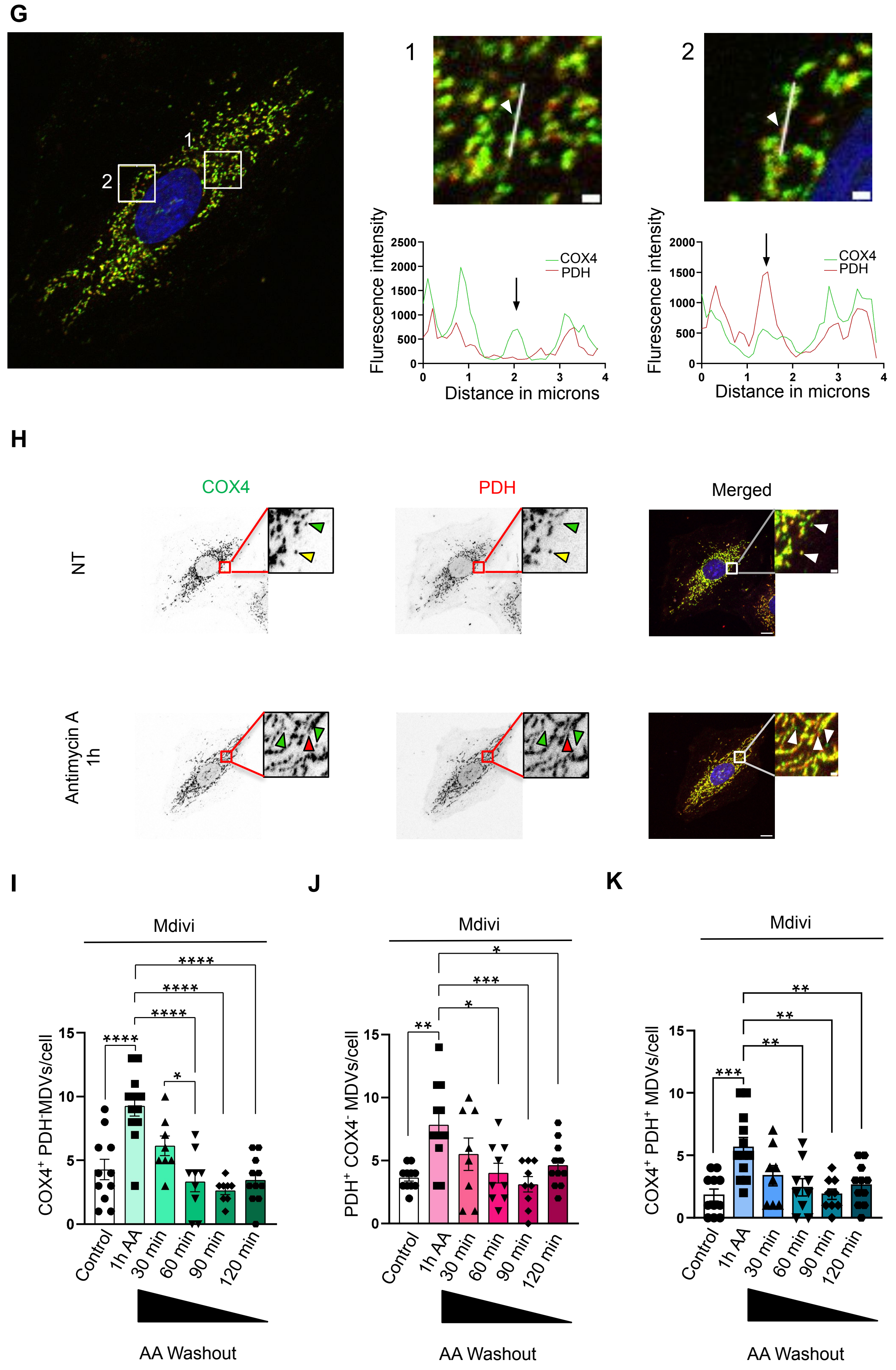

**Supplementary figure 1: Validation of MDVs in *in vitro* budding from cardiac mitochondria and in rat cardiomyoblasts.**

(A) Pie chart showing the mitochondrial localisation of the 110 mitochondrial proteins (present in at least two MS runs) comprising the MDV cargo. Details in Supplementary Data file S1.

(B) Pie chart showing the mitochondrial localisation of the 61 NEM sensitive high confidence MDV cargo (present in at least two MS runs). Details in Supplementary Data file S1.

(C) Reactome pathway analysis of the 110 mitochondrial proteins comprising MDV cargo. Shown are the top 10 pathways. Bars (purple) indicate the number of proteins of that pathway. Pathways covering more than 80% of the proteomic hits tend to be generic with little value and have been removed. The black line indicated the negative logarithm of the enrichment p-value. Details in Supplementary Data file S1.

(D) Reactome pathway analysis of the 61 NEM sensitive high confidence MDV cargo. Shown are the top 10 pathways. Bars (purple) indicate the number of proteins of that pathway. Pathways covering more than 80% of the proteomic hits tend to be generic with little value and have been removed. The black line indicated the negative logarithm of the enrichment p-value. Details in Supplementary Data file S1.

(E) Immunoblot of the budding reactions. The blot was probed for SDH and COX4. Control: budding reaction in the absence of N-ethylmaleimide (NEM). NEM: budding reaction in the presence of NEM (20mM). Mt. pellet: mitochondria pellet.

(F) Immunoblot of the sucrose density gradient separated vesicles. The blot was probed for SDH and COX4. Mt. pellet: mitochondria pellet.

(G) Validation of COX4$^+$PDH$^-$ MDVs and PDH$^+$ COX4$^-$ MDVs in H9C2 rat cardiomyoblasts. Line graphs to show COX4$^+$ PDH$^-$ MDVs(green) and PDH$^+$ COX4$^-$ MDVs (red) in region 1 and 2. Arrow on line graph points to the peak showing respective MDVs in region 1 and 2.

(H-K) Antimycin A (AA) washout assay (n=2, 8-12 cells). H9C2 rat cardiomyoblasts were pretreated with Mdivi (1μM), a DRP1 inhibitor for 1 hour prior to 1 hour treatment with AA (20μM). The AA stressor was removed at the end of 1 hour. The cells were fixed at 0min, 30 min, 60 min, 90 min and 120 min after removal of Antimycin A. (G)The cells were immunolabeled for COX4 (green) and PDH (red) and the nucleus was stained with DAPI (blue). Scale bar:10 μm (cell), 1 μm (inset). The green arrowheads indicate COX4$^+$ MDVs, red arrowheads indicate PDH$^+$ MDVs and yellow arrowheads indicate COX4$^+$ PDH$^+$ MDVs. (H) Quantification of COX4$^+$ MDVs. One-way ANOVA followed by Bonferroni post-hoc analysis, ****p-value< 0.0001, *p-value< 0.05. Error bars, SEM.(I) Quantification of PDH$^+$ MDVs. One-way ANOVA followed by Bonferroni post-hoc analysis, ***p-value< 0.001, **p-value< 0.01, *p-value< 0.05. Error bars, SEM. (J) Quantification of COX4$^+$ PDH$^+$ MDVs. One-way ANOVA followed by Bonferroni post-hoc analysis, ***p-value< 0.001, **p-value< 0.01. Error bars, SEM.

### Supplementary Figure 2

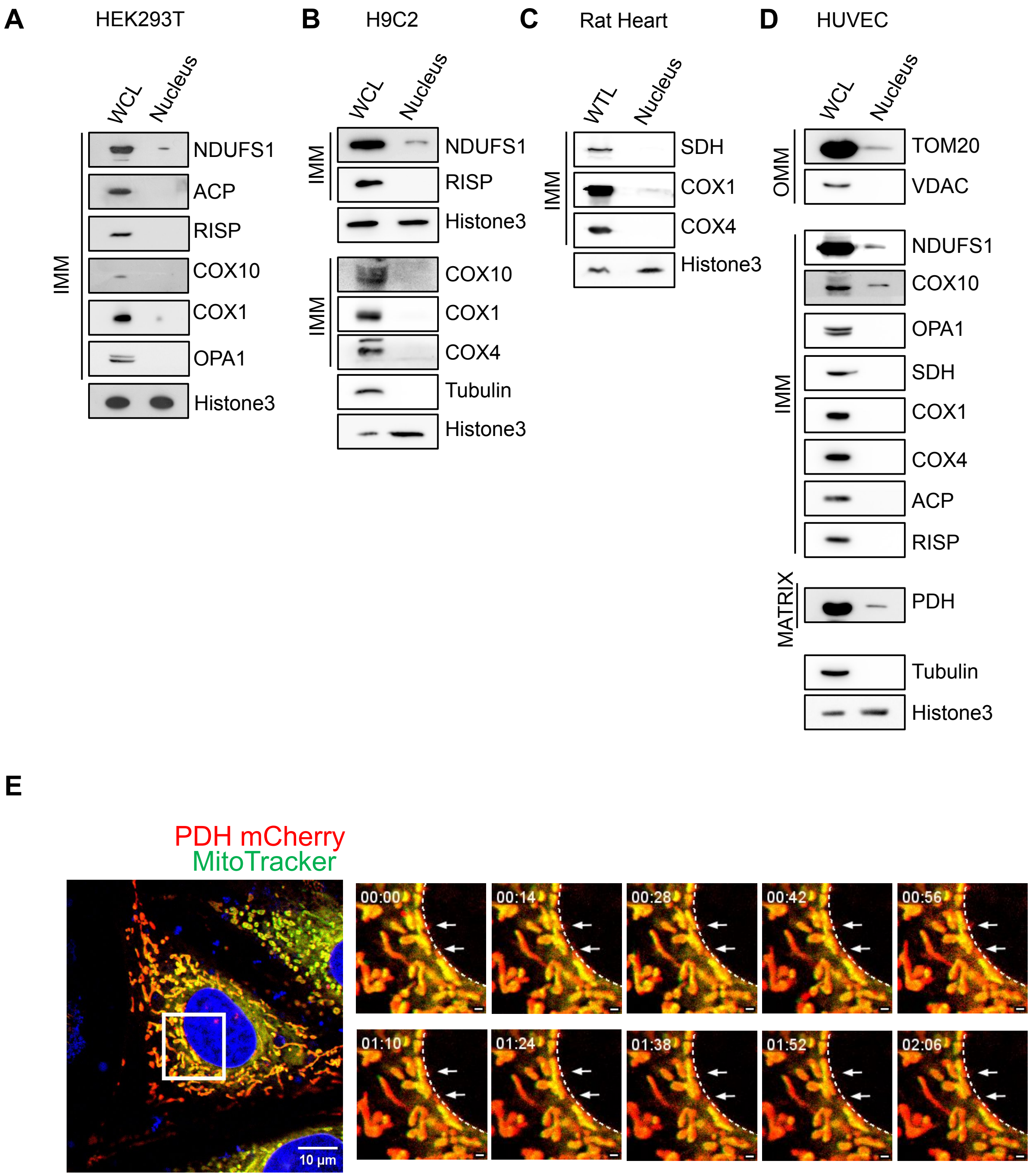

**Supplementary figure 2: Mitochondrial proteins transited to the nucleus**

(A) Immunoblot of the nuclei isolated from HEK293T. The blot was probed for the proteins as indicated. WCL: whole cell lysate.

(B) Immunoblot of the nuclei isolated from H9C2 cardiomyoblast. The blot was probed for the proteins as indicated. WCL: whole cell lysate.

(C) Immunoblot of the nuclei isolated from rat heart. The blot was probed for the proteins as indicated. WTL: whole tissue lysate.

(D) Immunoblot of the nuclei isolated from Human umbilical vein endothelial cells (HUVEC). The blot was probed for the proteins as indicated. WCL: whole cell lysate.

(E) Live imaging of H9C2 rat cardiomyoblasts overexpressing PDH mCherry (red). Mitochondria was stained by MitoTracker (500nM) pseudocolored in green and nucleus was stained with DAPI (blue).

### Supplementary Figure 3

A

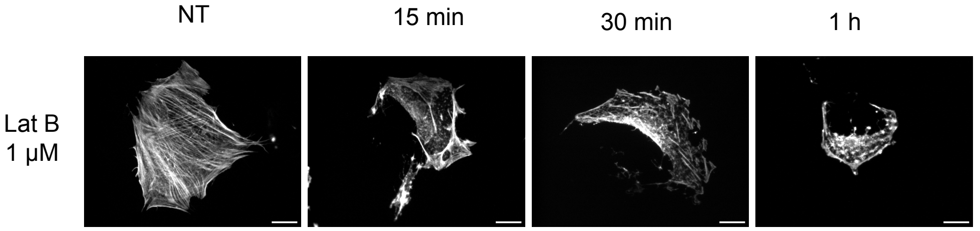


B

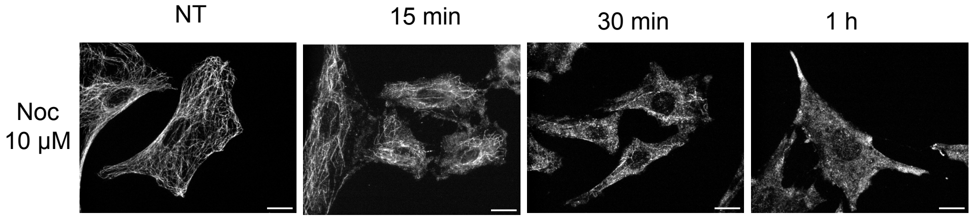


**Supplementary figure 3: Latrunculin B and nocodazole depolymerizes actin and tubulin respectively.**

(A) H9C2 cardiomyoblasts were treated with latrunculin B (1µM) and the cells were fixed with 100% methanol after 15 min, 30 min and 1 h of treatment. Actin was stained with phalloidin. Scale bar:10 µm.

(B) H9C2 cardiomyoblasts were treated with nocodazole (10µM) and the cells were fixed with 100% methanol after 15 min, 30 min and 1 h of treatment. Cells were immunolabeled for tubulin. Scale bar:10 µm.

### Supplementary Figure 4

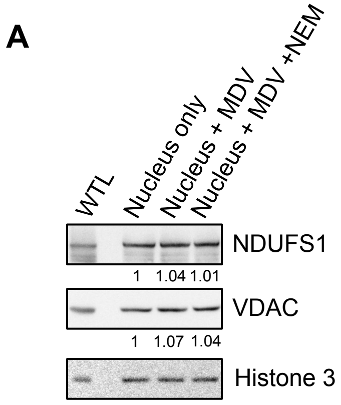


**Supplementary figure 4: Nuclear pools of NDUFS1 and VDAC are not NEM sensitive.**

(A) Immunoblot of the nuclei isolated after reconstitution as described in (Figure 4A). The blot was probed for NDUFS1, VDAC and Histone 3. On blot quantification with respect to Histone 3 is shown.

### Supplementary Figure 5

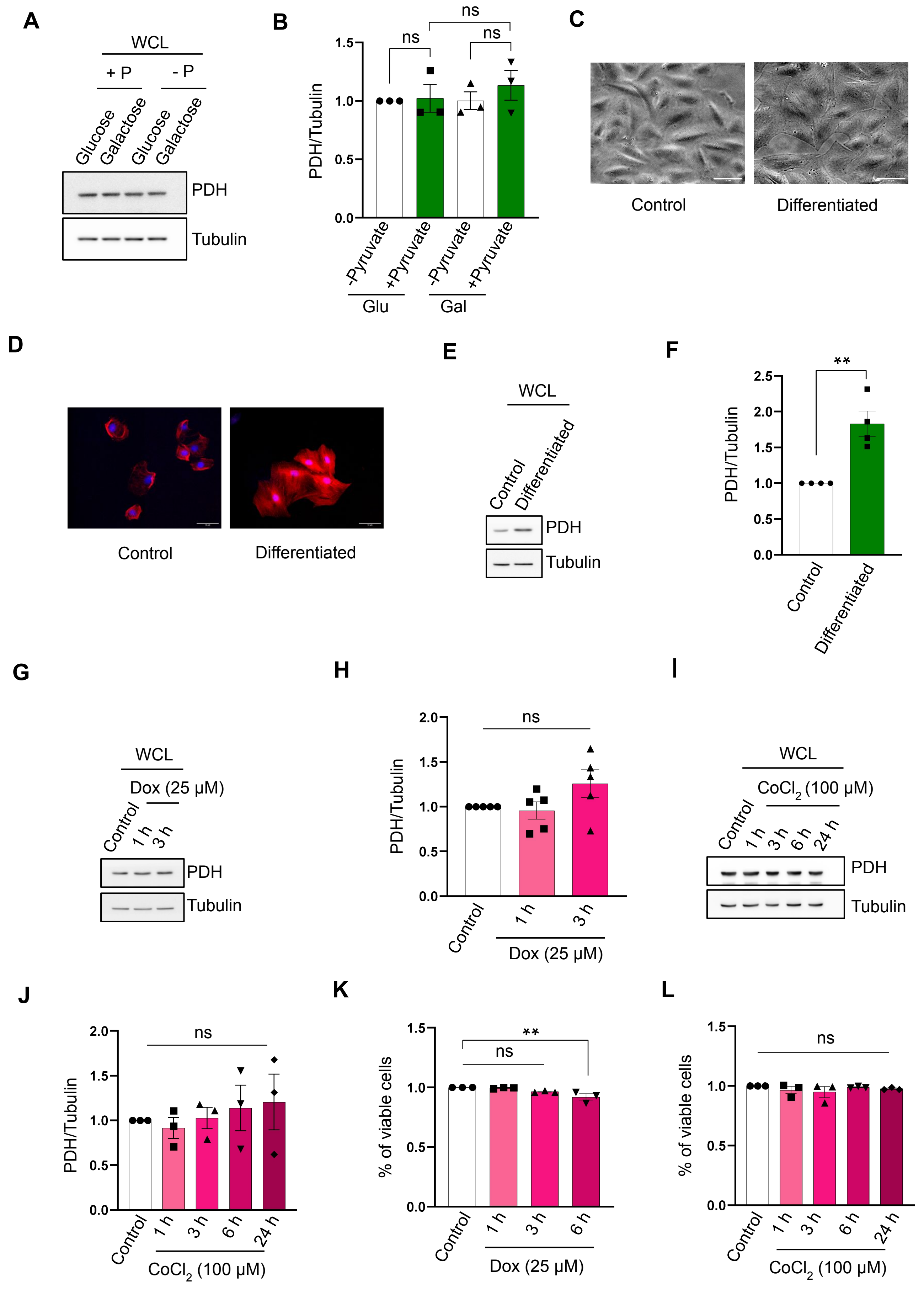

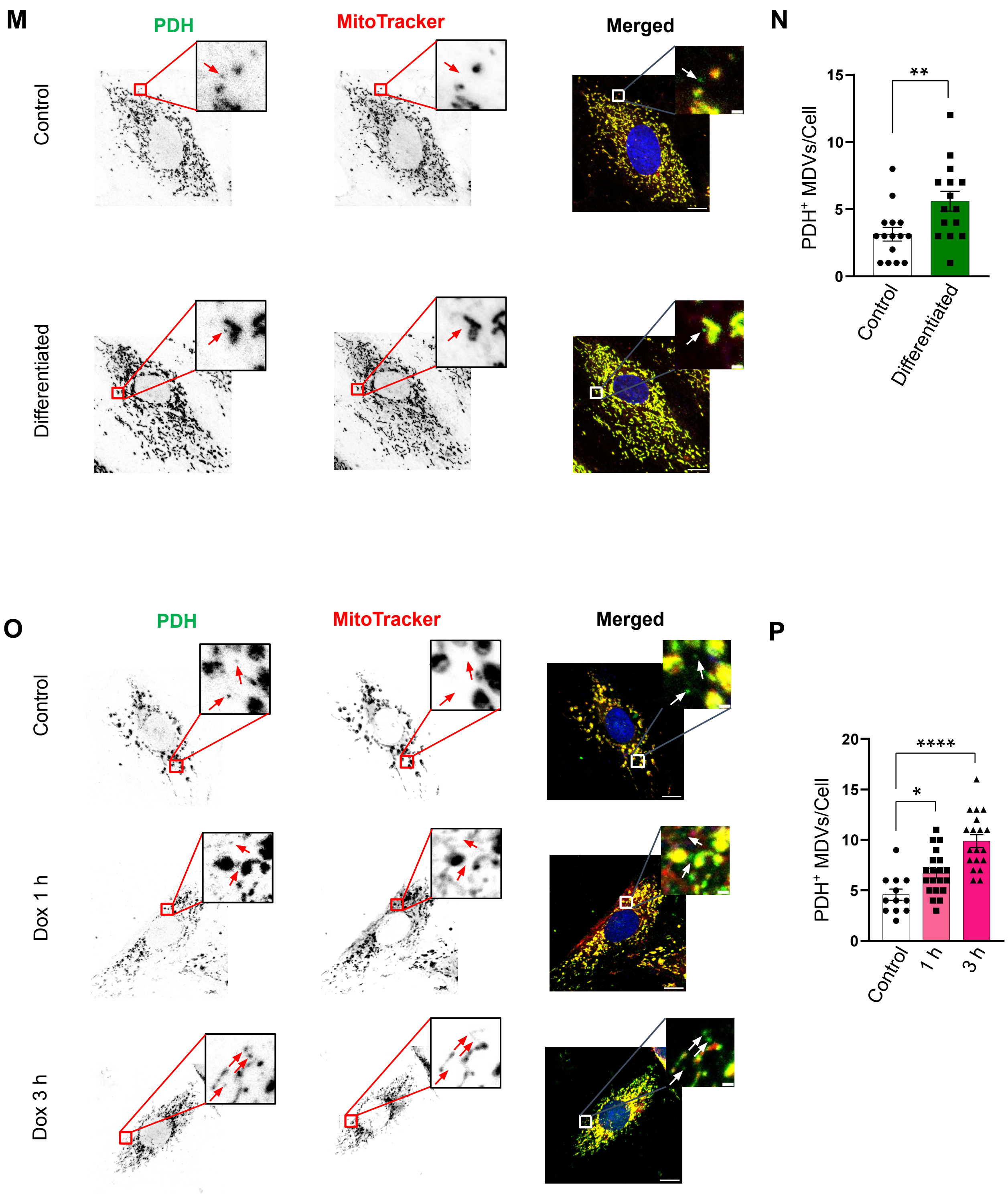
M
PDH
MitoTracker
Merged
Control
Differentiated
N
PDH+ MDVs/Cell
15
10
5
0
**
Control
Differentiated
O
PDH
MitoTracker
Merged
Control
Dox 1 h
Dox 3 h
P
PDH+ MDVs/Cell
20
15
10
5
0
****
*
Control
1 h
3 h

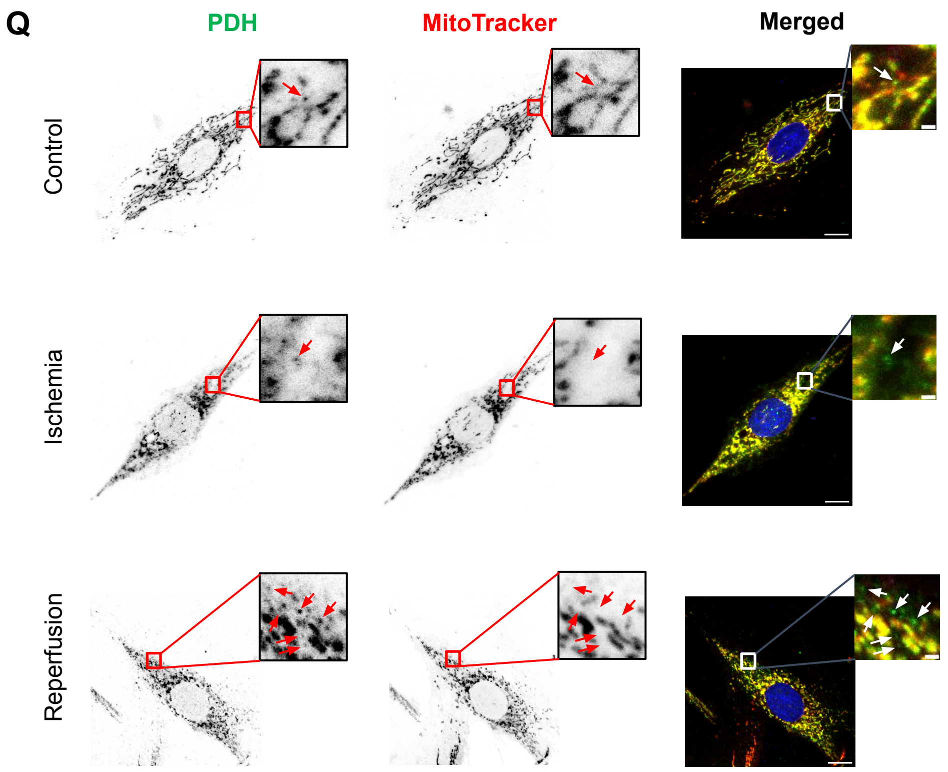

Q
PDH
MitoTracker
Merged
Control
Ischemia
Reperfusion


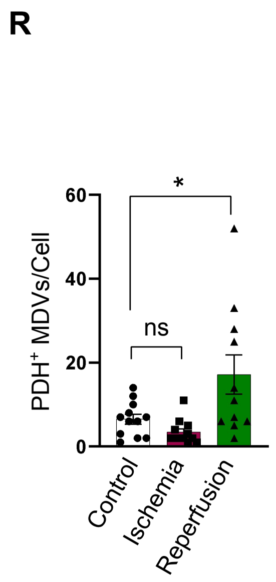

R
PDH+ MDVs/Cell
60
40
20
0
*
ns
Control
Ischemia
Reperfusion

**Supplementary Figure 5: Total PDH and PDH$^+$ MDV numbers in basal and pathophysiological conditions.**

(A,B) H9C2 cardiomyoblasts were cultured in 25 mM glucose and galactose-containing media (with a low 5 mM glucose) in the presence or absence of pyruvate (1 mM) for 24 hours. (A) Immunoblot of total cellular PDHE1, (B) Quantification of cellular PDH (n=3), PDH was normalized to tubulin. P: Pyruvate, Glu: Glucose, Gal: Galactose. RM one-way ANOVA followed by Bonferroni post-hoc analysis, ns: non-significant. Error bar, SEM.

(C-F) H9C2 cardiomyoblasts were differentiated into cardiomyocytes using retinoic acid (1 µM) for 5 days. (C) Widefield image of control and differentiated H9C2 cardiomyoblasts at Day 6, (D) Image of control and differentiated H9C2 cardiomyoblasts with Phalloidin staining. Scale bar: 10 µm. (E) Immunoblot of total cellular PDHE1, (B) Quantification of cellular PDH (n=4), PDH was normalized to tubulin. P: Pyruvate, Paired t-test,**p-value< 0.01. Error bar, SEM.

(G,H) H9C2 cardiomyoblasts were treated with Doxorubicin (25 µM) for 1 hour and 3 hours. (G) Immunoblot of total cellular PDHE1, (H) Quantification of cellular PDH (n=5), PDH was normalized to tubulin. One-way ANOVA followed by Bonferroni post-hoc analysis, ns: non-significant. Error bars, SEM.

(I,J) H9C2 cardiomyoblasts were treated with $CoCl_2$ (100 µM) to mimic hypoxia conditions. (I) Immunoblot of total cellular PDHE1, (J) Quantification of cellular PDH (n=3), PDH was normalized to tubulin. One-way ANOVA followed by Bonferroni post-hoc analysis, ns: non-significant. Error bars, SEM.

(K) Viability assay of H9C2 cardiomyoblasts under $CoCl_2$ (100 µM) using trypan blue dye (n=3). (L) Viability assay of H9C2 cells under doxorubicin (25 µM) treatment using trypan blue dye (n=3). One-way ANOVA followed by Bonferroni post-hoc analysis, **p-value< 0.01, ns: non-significant. Error bars, SEM.

(M) H9C2 cardiomyoblasts were differentiated into cardiomyocytes using retinoic acid (1 µM) for 5 days. Cells were immunolabeled for PDH E1-α (green); mitochondria and nucleus were stained with MitoTracker (red) and DAPI (blue) respectively. The arrows indicate PDH$^+$ MDVs (N) Quantification of PDH$^+$ MDVs (n=3, 5 cells/condition). Scale bar:10 µm (cell), 1 µm (inset). Unpaired t-test, **p-value< 0.01. Error bars, SEM.

(O) H9C2 cells were treated with doxorubicin (25µM) for 1 hour and 3 hours. The previously described staining protocol was followed. The arrows indicate PDH$^+$ MDVs (P) Quantification of PDH$^+$ MDVs (n=3, 5 cells/condition). Scale bar:10 µm (cell), 1 µm (inset). One-way ANOVA followed by Bonferroni post-hoc analysis, ****p-value< 0.0001, *p-value< 0.05. Error bars, SEM.

(Q) H9C2 cells were incubated in ischemia-reperfusion conditions as mentioned above. The previously described staining protocol was followed. The arrows indicate MDVs. (R) Quantification of PDH$^+$ MDVs (n=3, 5 cells/condition). Scale bar:10 µm (cell), 1 µm (inset). One-way ANOVA followed by Bonferroni post-hoc analysis, *p-value< 0.05. Error bars, SEM.
